# On the function of TRAP substrate-binding proteins: conformational variation of the sialic acid binding protein SiaP

**DOI:** 10.1101/2024.04.30.591957

**Authors:** Te-Rina J. King-Hudson, James S. Davies, Senwei Quan, Michael J. Currie, Zachary D. Tillett, Jack Copping, Santosh Panjikar, Rosmarie Friemann, Jane R. Allison, Rachel A. North, Renwick C.J. Dobson

## Abstract

Tripartite ATP-independent periplasmic (TRAP) transporters are analogous to ABC transporters in that they use a substrate-binding proteins to scavenge metabolites (*e.g.*, *N*-acetylneuraminate) and deliver them to the membrane components for import. TRAP substrate-binding proteins are thought to bind the substrate using a two-state (open and closed) induced-fit mechanism. We solved the structure of the TRAP *N*-acetylneuraminate substrate-binding protein from *Aggregatibacter actinomycetemcomitans* (*Aa*SiaP) in both the open ligand-free and closed liganded conformations. Surprisingly, we also observed an intermediate conformation, where *Aa*SiaP is mostly closed and is bound to a non-cognate ligand, acetate, which hints at how *N*-acetylneuraminate binding stabilises a fully closed state. *Aa*SiaP preferentially binds *N*-acetylneuraminate (*K*_D_ = 0.4 µM) compared to *N*-glycolylneuraminate (*K*_D_ = 4.4 µM), which is explained by the closed-*N*-acetylneuraminate bound structure. Small-angle X-ray scattering data alongside molecular dynamics simulations suggest the *Aa*SiaP adopts a more open state in solution than in crystal. However, the open unliganded conformation can also sample closed conformations. Molecular dynamics simulations also demonstrate the importance of water molecules for stabilising the closed conformation. Although our data is consistent with an induced fit model of binding, it is likely that the open unliganded conformation encompasses multiple states capable of binding substrate. The mechanism by which the ligand is released for import remains to be determined.

## Introduction

Sialic acids are an abundant family of nine-carbon sugars with many functions in mammals, including cell-cell recognition and signalling. More than 50 variants of sialic acid are known, but the most common in humans is *N*-acetylneuraminate (Neu5Ac). Sialic acids are scavenged by pathogenic and commensal bacteria as a source of energy or to evade the human immune system (1, 2). To utilise host-derived sialic acids, however, most bacteria must first import them using dedicated sialic acid transporters [(3) and reviewed in (4, 5)]. Disruption of bacterial sialic acid transporters (*i.e.*, knockouts) impairs the growth, survival, and colonisation of pathogenic bacteria (2, 6–9), highlighting the potential of these systems as antibacterial drug targets. Upon internalization, Neu5Ac is converted to *N-*acetylmannosamine, *N-*acetylmannosamine-6-phosphate, and *N-*acetylglucosamine-6-phosphate by the enzymes *N*-acetylneuraminate lyase (*nan*A), *N*-acetylneuraminate kinase (*nan*K), and *N*-acetylmannosamine-6-phosphate 2-epimerase (*nan*E) (10–14). Of particular interest are the sialic acid transporters of the tripartite ATP-independent periplasmic (TRAP) transporter family, as they are only present in bacteria and archaea (15) so could be targeted without affecting the human host.

TRAP transporters are secondary active transporters that transport specific ligands across bacterial membranes by co-transporting counter ions down an electrochemical gradient [(15–20) and recently reviewed here (21)]. Analogous to the ATP-binding cassette (ABC) transporter superfamily, TRAP transporters utilise a high-affinity substrate-binding protein that binds a target ligand with high affinity and delivers it to the membrane-spanning components for uptake (22, 23). In the sialic acid-specific TRAP system (SiaPQM), the substrate-binding protein, SiaP, is required for transport (18, 19, 21, 24) and associates with the membrane-spanning subunit, SiaQM, to facilitate translocation of sialic acid through an elevator-type transport mechanism.

Currently, 23 high-resolution structures are available for SiaP proteins (including various mutants) from just five species of Gram-negative bacteria: *Haemophilus influenzae, Vibrio cholerae, Fusobacterium nucleatum*, *Pasteurella multocida*, and *Photobacterium profundum* (listed in **Supplementary Table 1**). This wealth of structural data has provided insights into the molecular basis for the high affinity and specificity of SiaP proteins for sialic acids, particularly Neu5Ac and *N*-glycolylneuraminate (Neu5Gc) (25–28). The structural core of substrate-binding proteins consists of two globular domains connected by a variable hinge region (29), which is formed by two β-strands and a unique extended α-helix in SiaP. Ligands bind within the cleft between the two domains, which close around the bound ligand *via* a hinge-bending conformational change often described as a ‘Venus flytrap’ mechanism (30, 31). The available crystal structures for SiaP without ligand (apo/unbound state) adopt open conformations, suggesting that closure is strictly ligand-induced, consistent with a two-state induced fit binding model (27).

One uncertainty is the extent to which unbound SiaP undergoes intrinsic closure in solution like other ABC substrate-binding proteins (32–35). SiaP conformational dynamics have been investigated using pulsed electron-electron double resonance spectroscopy and single molecule FRET, which support the proposal that *V. cholerae* SiaP (*Vc*SiaP) exclusively occupies an open conformation in the absence of ligand and that ligand binding induces and stabilises closure (36, 37). Molecular dynamics simulations uniquely developed to reproduce the DEER signal from these experiments suggest that *Vc*SiaP generally adopts a more open conformation in solution than the observed from the crystal structure, and is conformationally flexible, but does not fully close without the ligand (38). Together, these data present a sensible model for SiaP dynamics, although it is unclear whether this model can be applied to all SiaP proteins, or whether there is variation between homologs. Furthermore, the precise details of how sialic acid binding may trigger the conformational change in SiaP are yet to be elucidated.

Here, we report the functional, biophysical, and structural characterisation of the substrate-binding protein of the sialic acid TRAP transporter from *Aggregatibacter actinomycetemcomitans* (*Aa*SiaP), a known periodontal pathogen that causes infective endocarditis (39, 40). We demonstrate that *Aa*SiaP preferentially binds Neu5Ac over Neu5Gc with nanomolar affinity and that sodium ions are not involved in binding. Our findings suggest that *Aa*SiaP can adopt a mostly closed conformation that is stabilised by acetate, a small non-cognate ligand. Molecular dynamics simulations and small angle X-ray scattering data support the notion that *Aa*SiaP has greater conformational flexibility than is evident from the crystal structures.

## Results

### Bioinformatic analysis demonstrates that AaSiaP shares key residues for Neu5Ac binding

We first conducted a bioinformatic analysis of the putative *siaP* gene from *A. actinomycetemcomitans* (*Aa*SiaP) for two reasons. Firstly, to verify that *siaP* is correctly annotated in the *A. actinomycetemcomitans* genome, a basic local alignment search tool (BLAST) analysis was performed with the *Aa*SiaP amino acid sequence. The *Aa*SiaP sequence showed a high level of similarity to the amino acid sequences of confirmed sialic acid substrate-binding proteins (SiaP), consistent with its proposed role in sialic acid metabolism. A sequence alignment (**Figure 1**) demonstrates that many of the key residues for sialic acid binding and specificity [including R127, R147, F170, and H209 (26, 28)] are conserved in the *Aa*SiaP primary sequence, supporting its identity as a sialic acid substrate-binding protein. Interestingly, the well-conserved surface residue N150, which could be important for transport, was also present in *Aa*SiaP and 11 of 12 homologs but substituted to glycine in *Fn*SiaP. Secondly, an analysis of the mechanism of ligand binding for *Aa*SiaP requires a known protein structure. The bioinformatic survey provided 12 sequences of homologous proteins (50–95% sequence similarity, **Supplementary Figure 1**), with four of these structurally characterised using X-ray crystallography, which can be used as templates for molecular replacement of *Aa*SiaP (**Figure 1**).

**Figure 1.**
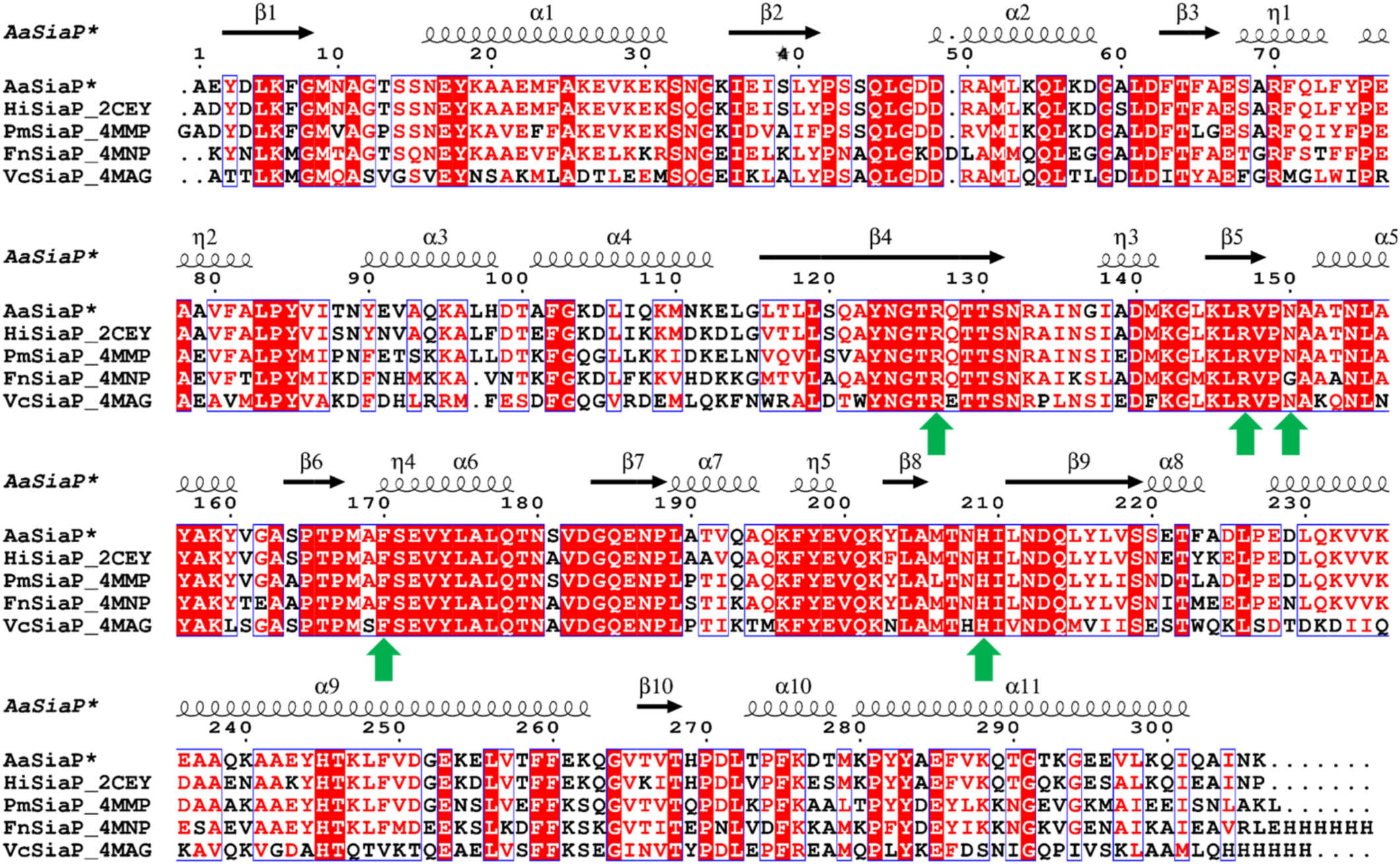
Multiple sequence alignment of Neu5Ac-binding proteins homologous to *Aa*SiaP with structures deposited in the PDB. The residue numbering and secondary structure is based on *Hi*SiaP. Sequence identity between *Aa*SiaP each homologue is as follows: *Hi*SiaP 88.5%, *Pm*SiaP 75.2%, *Fn*SiaP 66.5%, and *Vc*SiaP 51.6%. Red background indicates complete sequence identity and red letters indicate residues with similar physicochemical properties. *Recombinant *Aa*SiaP used in this study had an N-terminal cleavage fragment comprising an M and D residue scar not shown here. Created in ESPript 3.0 (https://espript.ibcp.fr/ESPript/cgi-bin/ESPript.cgi). A broader sequence alignment of Neu5Ac-binding proteins is in **Supplementary Figure 1**. Residues discussed here are marked with a green arrow.

### AaSiaP demonstrates a preference for N-acetylneuraminate over N-glycolylneuraminate

*Aa*SiaP was recombinantly expressed in *Escherichia coli* for subsequent structural and biophysical analyses. To facilitate folding of the protein in the *E. coli* periplasm, the native *A. actinomycetemcomitans* periplasmic signal sequence was replaced with the *pelB* periplasmic signal sequence. Recombinant *Aa*SiaP was highly expressed and could be isolated from the periplasmic fraction (**Supplementary Figure 2A**) and purified by anion-exchange, hydrophobic-interaction, and size-exclusion chromatography (**Supplementary Figure 2B**). *Aa*SiaP eluted from the size-exclusion column as a single peak (**Supplementary Figure 2C**), suggesting that it is a monodisperse species and remains stable throughout purification.

To confirm the function of *Aa*SiaP we used two solution-based techniques that detect temperature-related changes in fluorescent intensity to estimate the affinity and thermal stabilisation of ligand binding (**Table 1**).

**Table 1.** Estimated dissociation constant (*K*_D_) values for *Aa*SiaP binding to Neu5Ac and Neu5Gc from microscale thermophoresis and thermal stability of Neu5Ac binding from differential scanning fluorimetry in the presence of either K^+^ or Na^+^.

| Dissociation constant (95% CI) <sup>a</sup> |  |  |
| --- | --- | --- |
| Ligand | *K*<sub>D</sub> *Aa*SiaP-FITC (nM) |  |
| Neu5Ac | 430 (136–724) |  |
| Neu5Gc | 4400 (2600–6200) |  |
| Thermal stability ± SD <sup>b</sup> |  |  |
| Salt | *Aa*SiaP *T*<sub>m</sub> (°C) | *T*<sub>m</sub> + 50 μM Neu5Ac (Δ*T*<sub>m</sub>) (°C) |
| NaCl | 50.7 ± 0.6 | 57.2 ± 0.9 (+ 6.4) |
| KCl | 51.1 ± 0.3 | 58.6 ± 0.6 (+ 7.5) |

<sup>a</sup> Determined by microscale thermophoresis.
<sup>b</sup> Determined by differential scanning fluorimetry.
\* *Aa*SiaP in Tris-NaCl (+) and Tris-KCl (-).

Microscale thermophoresis was used to estimate the binding affinity of *Aa*SiaP to the sialic acids *N*-acetylneuraminate (Neu5Ac) and *N*-glycolylneuraminate (Neu5Gc) (**Figure 2A**, **Table 1**). The protein was labelled with fluorescein-5-isothiocyanate (FITC) directed towards the N-terminal free amine (41) located in the less dynamic domain I to minimise potential fluorophore-protein interactions. However, this conservative labelling approach gave a low fluorophore/protein molar ratio and some fluorescent counts were below the recommended threshold due to rapid photobleaching (**Supplementary Figure 3B-D**). Despite this, we obtained estimates of *Aa*SiaP binding affinity (*K*_D_) of 430 nM for Neu5Ac (95% CI: 136–724 nM) and 4400 nM for Neu5Gc (95% CI: 2600–6200 nM) that are in good agreement with published values for SiaP homologues from *H. influenzae, P. multocida,* and *F. nucleatum* (**Supplementary Table 1**).

**Figure 2.**
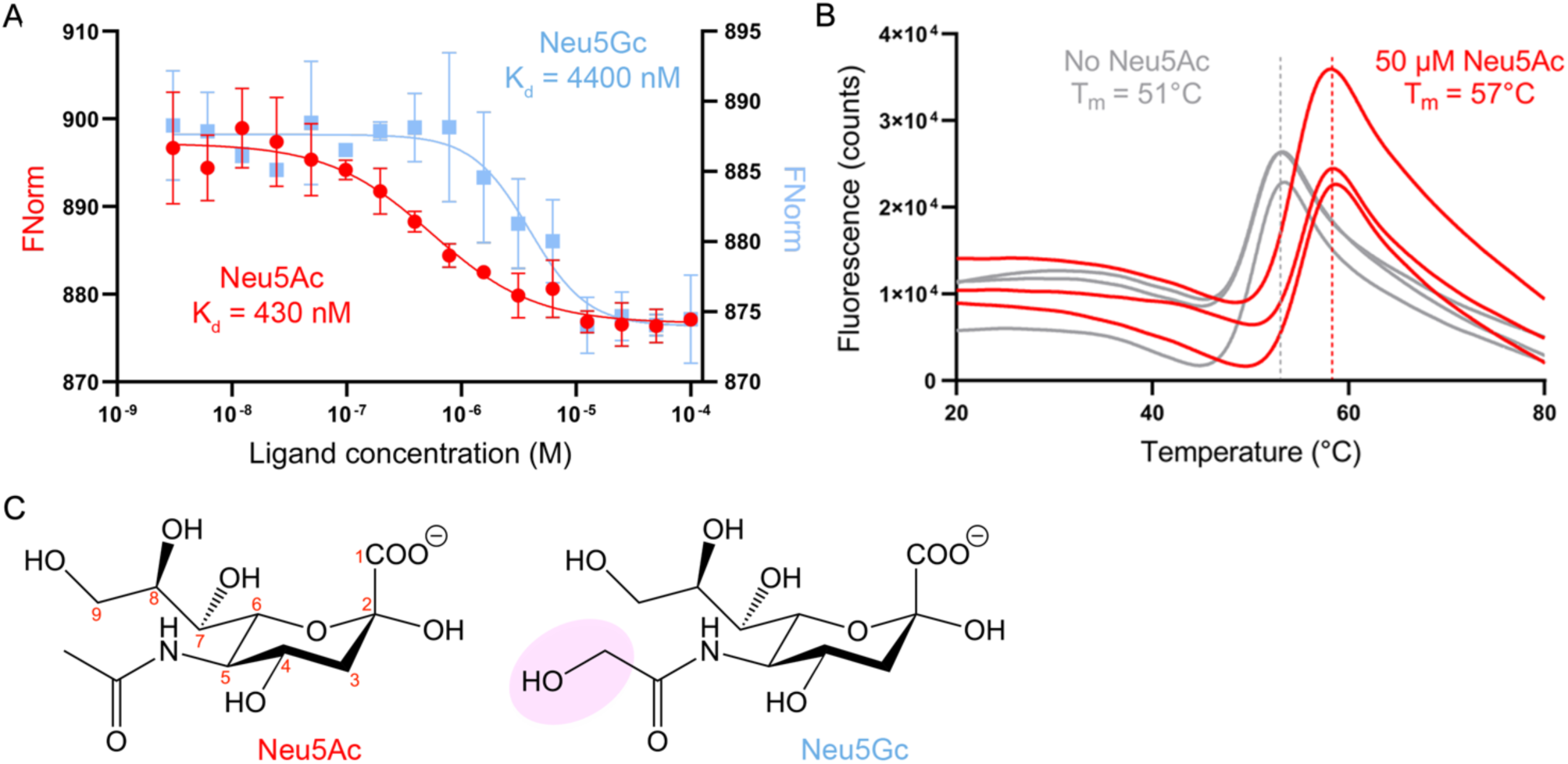
Ligand binding to *Aa*SiaP. **A**) *Aa*SiaP (75 nM) binding curve and affinity (*K*_D_) for Neu5Ac (red circle) and Neu5Gc (light blue squares) determined by microscale thermophoresis, errors bars indicate standard deviation. **B**) Thermal shift assay results for *Aa*SiaP (1 µM) showing increase in melting temperature (T_m_) from apo (grey curves) with addition of 50 µM Neu5Ac (red curve), determined by differential scanning fluorimetry, individual replicates are shown. **C**) The structures of Neu5Ac and Neu5Gc, highlighting their difference (shaded pink).

Neu5Ac binding was also estimated using differential scanning fluorimetry thermal shift assays with unlabelled *Aa*SiaP (**Figure 2B**, **Table 1**). This experiment demonstrates that *Aa*SiaP is stable at room temperature (T_m_ with Tris-NaCl = 50.7 ± 0.6 °C) and is further stabilised by Neu5Ac binding (T_m_ with 50 µM Neu5Ac = 57.2 ± 0.9 °C [an increase of 6.4 °C]) to a similar extent as *Hi*SiaP (25). TRAP transporters use Na^+^ ion gradients to drive transport (17, 19, 21, 24) and one intriguing possibility is that the Na^+^ ions are also bound and delivered to the transporter by the substrate-binding protein. *Aa*SiaP was similarly stabilised by Neu5Ac in the absence of Na^+^ ions (T_m_ with Tris-KCl + 50 µM Neu5Ac = 58.6 ± 0.6 °C [an increase of 7.5 °C]). Titration of Neu5Ac into *Aa*SiaP and monitoring for a change in melting temperature estimated the *K*_D_ to be 54.6 μM (95% CI: 21–122 μM) when in Tris-NaCl buffer, and this is largely unchanged in the absence of Na^+^ ions (Tris-KCl buffer), where the *K*_D_ is estimated to be 37.9 μM (95% CI: 23–62 μM) (**Supplementary Figure 3A**). The discrepancy in the *K*_D_ between the MST and DSF experiments is expected because in DSF the temperature is increased.

Together, microscale thermophoresis and thermal shift assays demonstrate that *Aa*SiaP binds both Neu5Ac and Neu5Gc, but with a clear preference for Neu5Ac. Sodium ions (used by TRAP transporters for symport) do not appear to be directly involved in the binding of Neu5Ac to *Aa*SiaP—consistent with available crystal structures that do not show any obvious sodium binding sites.

### The crystal structures of apo- and Neu5Ac-bound AaSiaP reveal a series of distinct conformations

To better define the molecular mechanism by which *Aa*SiaP binds Neu5Ac and to complement our solution data, we determined the crystal structure of *Aa*SiaP in both the absence and presence of Neu5Ac. Of the 23 SiaP structures deposited in the Protein Data Bank (PDB), 16 are variations of *Hi*SiaP (residue substitutions, different ligands) and another four are variations of *Vc*SiaP (**Supplementary Table 2**), so there is a need to study further homologues to understand the generality of the binding mechanism. Moreover, there are just two structures of SiaP in the open conformation (*i.e.*, apo-SiaP), limiting a generalised understanding of how these proteins bind Neu5Ac with high affinity.

Apo-*Aa*SiaP easily afforded crystals in a variety of conditions, the best of which diffracted to a maximum resolution of 2.45 Å. Despite having very good search models, we initially struggled to solve the phases by molecular replacement for the apo-*Aa*SiaP structure. The Matthew’s coefficient estimated four to six molecules in the asymmetric unit, but molecular replacement phasing attempts using an unliganded *Hi*SiaP structure in the open conformation as the search model (89% sequence identity, PDB ID: 2CEY) placed only two molecules. Attempts to phase with Neu5Ac-bound *Pm*SiaP in the closed conformation (75% sequence similarity, PDB ID: 4MMP) also only yielded a partial solution of two molecules. A successful solution was found by searching for two molecules of *Hi*SiaP structure in the open conformation and two *Pm*SiaP monomers in a closed conformation, despite no Neu5Ac being added during purification or crystallisation. Thus, the crystal structure revealed that apo-*Aa*SiaP is present in at least two distinct conformations (**Figure 3A**). The final refined model had an *R_free_* of 24.6% and a *R_work_* of 18.5%, with reasonable geometry and Ramachandran outliers for the resolution (2.45 Å), as judged by MolProbity (42, 43) (**Table 2**).

**Figure 3.**
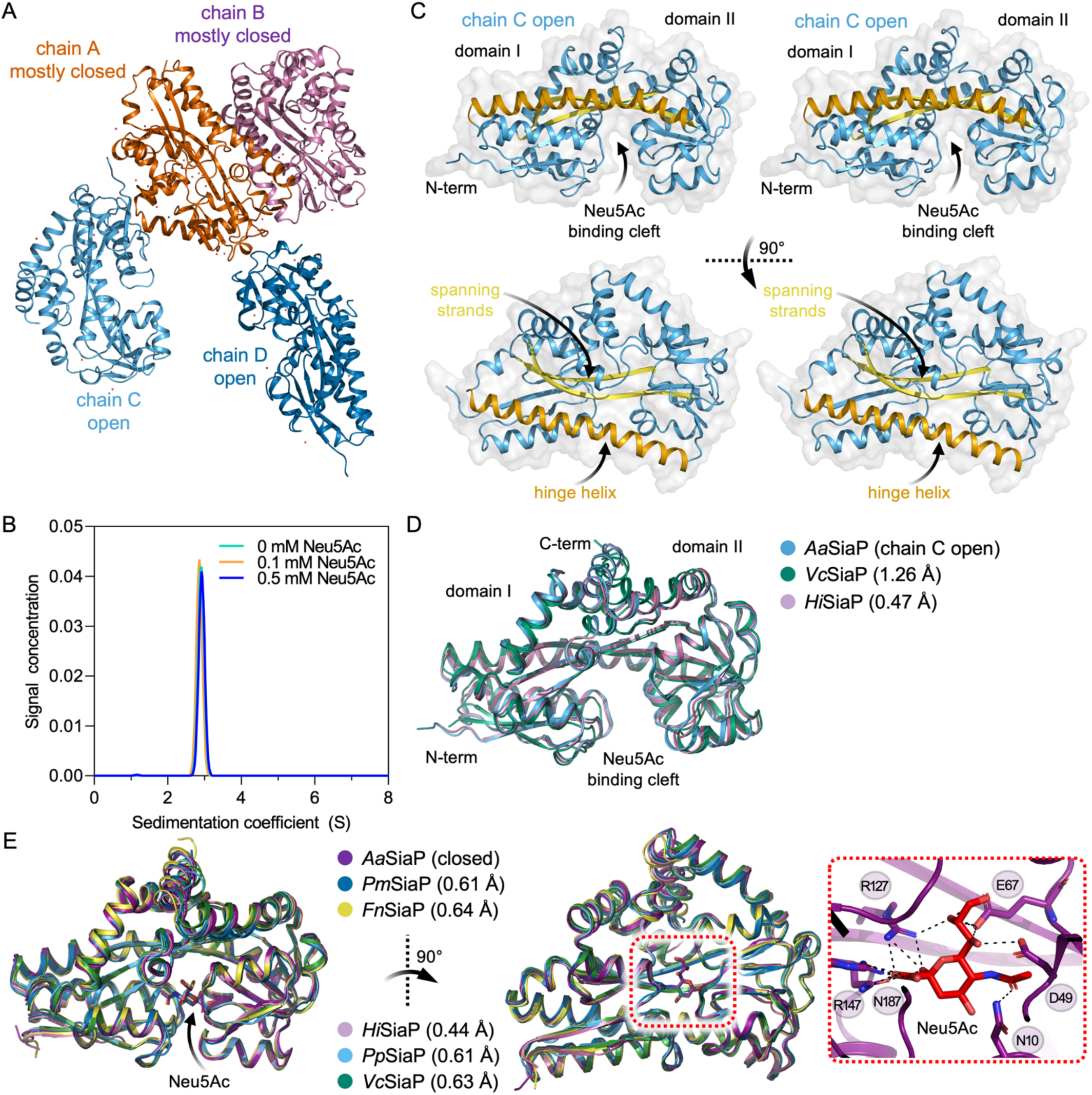
Crystal structure of *Aa*SiaP. **A**) Asymmetric unit of ligand-free *Aa*SiaP contains fours monomers. Chains A and B are in a mostly closed conformation, whereas chains C and D are in an open conformation. PISA analysis (https://www.ebi.ac.uk/pdbe/pisa/) demonstrates that the largest interface is between chains A and B (~530 Å^2^). **B**) Sedimentation velocity AUC analysis of *Aa*SiaP at various concentrations of Neu5Acshows a well-defined single peak at ~2.8 S. This is indicative of a monodisperse and monomeric protein in solution that is not affected by the presence of Neu5Ac. Fits to the data are in **Supplementary Figure 4**. **C**) Stereo plots (cross eyed) showing the overall architecture of *Aa*SiaP (chain C). **D**) An overlay of homologous SiaP structures in the equivalent open conformation [*Vc*SiaP (4MAG (28)) and *Hi*SiaP (2CEY (27))] shows that the structure is highly conserved. **E**) An overlay of homologous SiaP structures in the equivalent ligand-bound, closed conformation [*Pm*SiaP (4MMP (28)), *Fn*SiaP (4MNP (28)), *Hi*SiaP (6H76 (25)), *Pp*SiaP (7T3E (17)), and *Vc*SiaP (7A5Q (37))]. In all structures Neu5Ac is bound in the same pose. The inset shows the *Aa*SiaP residues binding Neu5Ac.

**Table 2.** Crystallographic data collection and refinement statistics for recombinant *Aa*SiaP.

|  | **Apo-*Aa*SiaP** | **Neu5Ac-bound *Aa*SiaP** |
| --- | --- | --- |
| **data collection** |  |  |
| temperature (K) | 100 | 100 |
| detector | EIGER X 16M (Dectris) | EIGER X 16M (Dectris) |
| crystal-to-detector distance (mm) | 300 | 200 |
| space group | *C* 2 | *P* 2<sub>1</sub> |
| unit cell parameters [a, b, c (Å); β (°)] | 138.7, 141.4, 109.4; 112.5 | 66.2, 49.5, 121.4; 92.8 |
| resolution range (Å) | 48.64–2.58 (2.65–2.58) | 45.86–1.90 (2.0–1.90) |
| no. of unique reflections | 61,226 (2,476) | 62,386 (6,175) |
| multiplicity (outer shell) | 3.6 (3.7) | 2.0 (2.0) |
| completeness (%) (outer shell) | 100.0 (99.9) | 99.76 (99.72) |
| *I*/σ(*I*) (outer shell) | 6.3 (1.6) | 6.2 (1.18) |
| CC<sub>(1/2)</sub> (outer shell) | 0.995 (0.783) | 0.995 (0.505) |
| *R*<sub>merge</sub> (outer shell) | 0.087 (0.612) | 0.064 (0.641) |
| **refinement statistics** |  |  |
| no. of reflections used | 58,058 | 62,286 |
| no. of protein atoms | 9,609 | 4,862 |
| no. of water molecules | 49 | 487 |
| *R*<sub>work</sub> (%) | 18.9 | 20.0 |
| *R*<sub>free</sub> (%) | 22.0 | 23.1 |
| reflections used for *R*<sub>free</sub> | 3,158 (243) | 3,107 (127) |
| Ramachandran (favoured/allowed/disallowed) | 99.6, 0.3, 0.08 | 98.4, 1.6, 0 |
| r.m.s.d. bond lengths (Å) | 0.015 | 0.014 |
| r.m.s.d. angles (°) | 1.58 | 1.461 |
| clashscore (*Phenix*) | 4.4 | 3.16 |
| PDB id | 9BH3 | 9BHF |

We also solved a 1.9 Å crystal structure of Neu5Ac-bound *Aa*SiaP. Molecular replacement was straightforward in this case using a liganded *Hi*SiaP structure (PDB ID: 2WYK). The structure closely resembles the Neu5Ac-bound *Hi*SiaP structure (root mean square deviation (r.m.s.d.) = 0.44 Å, across 283 C_α_-atoms) and can be described as a fully closed, bound conformation. For both models, refinement was initially performed with simulated annealing to remove model bias.

PISA analysis (44) demonstrates that the most significant interface is between the mostly closed monomers (chains A and B, **Figure 3A**) in the apo-*Aa*SiaP crystal (~530 Å^2^). To check whether *Aa*SiaP can oligomerise we conducted analytical ultracentrifugation sedimentation velocity experiments in the presence and absence of Neu5Ac (**Figure 3B**). In all cases, the samples present as a single symmetrical peak at ~2.9 S. The estimated mass of *Aa*SiaP is 34–35 kDa based on these experiments, which is consistent with the mass calculated from the amino acid sequence (34.3 kDa). No detectable change in the frictional ratio (a measure of asymmetry) was found when Neu5Ac was added (*f*/*f*_o_ = 1.27–1.28), suggesting that sedimentation velocity experiments are not sensitive enough to detect the change in shape.

The monomeric structure of *Aa*SiaP is similar to other reported SiaP structures (see references in **Supplementary Table 2**), first described for *Hi*SiaP (27). The structure has two domains that comprise residues from both the N-terminal and the C-terminal ends of the sequence (**Figure 3C**). The domains are bridged by two long β-strands (spanning strands) and a long α-helix (the hinge helix) and form a cleft within which Neu5Ac binds.

The observation that of the 23 SiaP structures deposited in the PDB, just two structures are in the open, ligand free conformation (4MAG and 2CEY, **Supplementary Table 2**) may reflect some difficulty in crystallising SiaP proteins in the absence of Neu5Ac due to conformational flexibility. Anecdotally, we found *Aa*SiaP more easily crystallised in the presence of Neu5Ac, and the resolution is considerably better, which is consistent with the large stabilising effect of Neu5Ac (T_m_ + 50 µM Neu5Ac = 57.2 ± 0.9 °C, an increase of 6.4 °C).

Comparing the open conformation with equivalent homologues in the PDB shows that the open structure is highly conserved (**Figure 3D**), *Aa*SiaP to *Vc*SiaP (4MAG) r.m.s.d. = ~1.23 Å across 278 C_α_-atoms and *Aa*SiaP to *Hi*SiaP (2CEY) r.m.s.d. = 0.5 Å across 279 C_α_-atoms (27, 28) (**Supplementary Table 3**). The closed Neu5Ac-bound structure is also highly conserved when compared with equivalent homologues (r.m.s.d. = 0.4–0.6 Å across five homologues, **Figure 3E**). In all available cases the binding pose for Neu5Ac is identical and the residues that bind Neu5Ac are well conserved (**Supplementary Figure 5**). Recent work has highlighted the importance of water networks for Neu5Ac binding (25) and again we find that the water network around Neu5Ac is also highly conserved (**Supplementary Figure 5**).

### Conformational change upon Neu5Ac binding

We compared the three structures of *Aa*SiaP to define the motions the protein undergoes in solution and when binding Neu5Ac. An overlay of the open and mostly closed conformations of apo-*Aa*SiaP (**Figure 4A,B**) differed by an r.m.s.d. of ~1.5 Å across 283 C_α_-atoms (**Supplementary Table 2**). The two mostly closed conformations more closely resemble the Neu5Ac-bound *Aa*SiaP (chain A r.m.s.d. = 0.4 Å, chain B = 0.3 Å) and *Hi*SiaP structures (r.m.s.d. = ~0.4 Å; **Supplementary Table 2**). The mostly closed unbound structure represents a unique intermediate conformation that, to our knowledge, has not been reported for TRAP substrate-binding proteins but has been observed in ABC substrate-binding proteins (32, 35).

**Figure 4.**
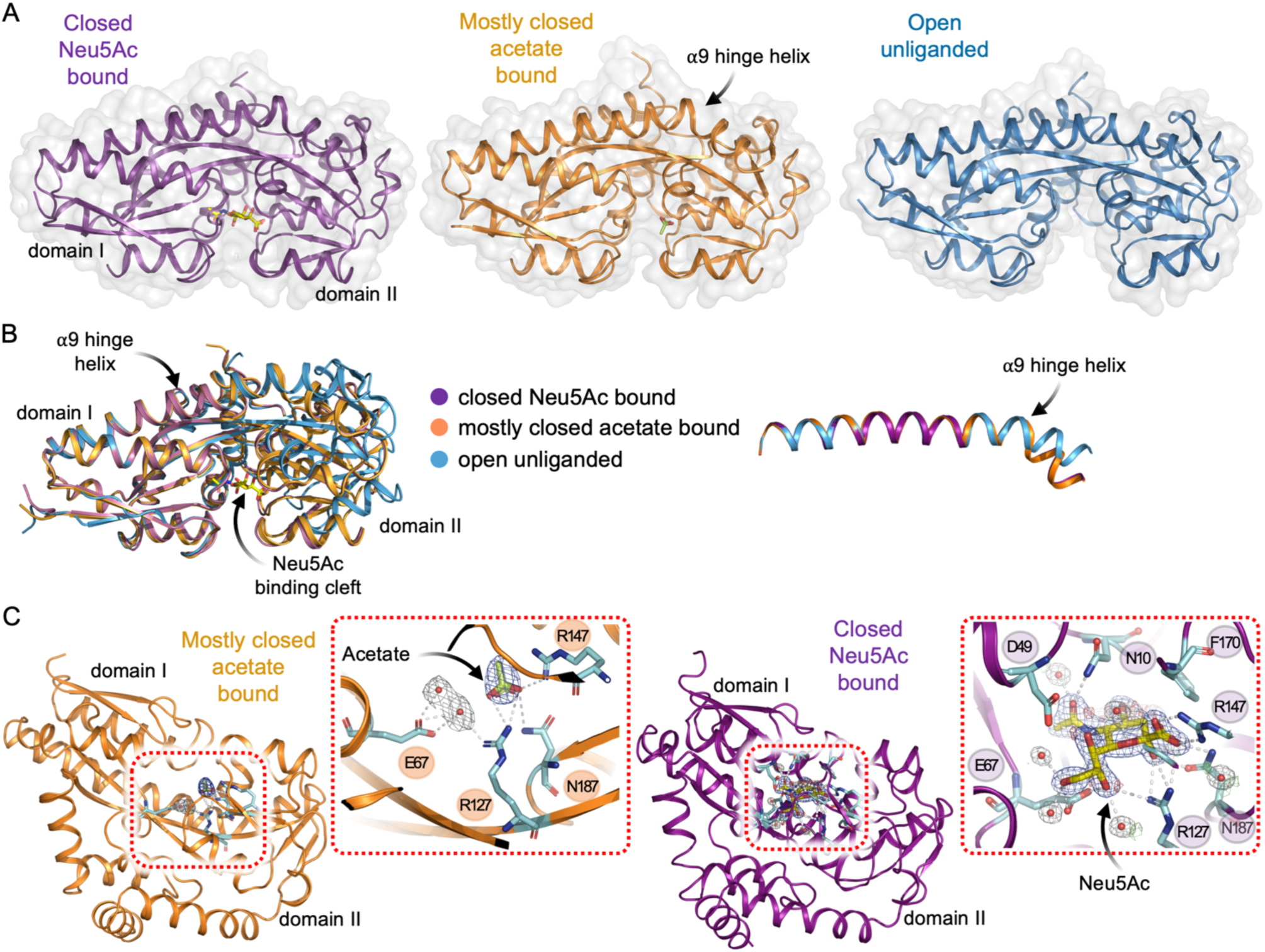
| The conformational landscape of *Aa*SiaP. **A)** In purple, the closed Neu5Ac-bound conformation, showing Neu5Ac bound deep within a cleft between the N-terminal (domain I) and C-terminal (domain II) lobes. In orange is the mostly closed conformation, where acetate is also bound by twin conserved arginine residues of domain II. In blue, the open unliganded conformer from the same crystal structure as the mostly closed conformer. As depicted, the α9 hinge helix running between the domains undergoes a substantial hinge bending motion to allow the two domains to come together. **B**) An overlay of the three conformations of *Aa*SiaP reported here. The bend in the α9 hinge helix is highlighted in the right pane. **C**) Zoom-in of the substrate-binding site, highlighting ordered water molecules and hydrogen bonding interactions (white dashes). Electron density (2mF_o_-DF_c_ contoured at 1σ, mF_o_-F_c_ contoured at 3σ) at the substrate-binding site of the the mostly closed acetate-bound structure (left) and closed Neu5Ac-bound structure (right).

The closed conformation of TRAP substrate-binding proteins has only been observed in the presence of substrate, so we checked the mostly closed structure for a bound ligand. An electron density peak larger than that of a water molecule was evident near the highly conserved R147 and R127 residues that would otherwise coordinate the carboxyl group of Neu5Ac (**Figure 4C**). This density is consistent with acetate, a small organic anion with a carboxyl moiety that was present in both crystallisation conditions. Acetate makes favourable interactions with residues R127, R147 and N189 on domain II within the binding site of *Aa*SiaP, effectively mimicking the carboxyl moiety of Neu5Ac. However, unlike Neu5Ac, acetate does not bridge the two domains which fails to explain the closure of the domains. We note that in chain B of the mostly closed structure, acetate also makes contact with a chain of water molecules (**Figure 4C**), which in turn contacts E67 in domain I, a residue that normally interacts with Neu5Ac. Although the resolution of the apo-*Aa*SiaP structure (2.58 Å) is lower compared to that of the Neu5Ac structure (1.9 Å), resulting in fewer evident waters. Furthermore neither the mostly closed nor open structures exhibit the shell of highly ordered and conserved waters observed in the Neu5Ac binding site when Neu5Ac is present. It remains unclear whether acetate binding induces a partial closure of *Aa*SiaP. However, if it does, this would suggest that the twin arginine residues to which acetate binds, and perhaps also E67 from domain I, are involved in sensing the substrate and potentially play a role in triggering or stabilizing the conformational change towards partial closure, as originally proposed by (28).

The change from the open to the closed conformation in *Aa*SiaP induces a significant kink in the hinge helix α9 (**Figure 4B**), as reported in other SiaP proteins. This helix has been proposed to function as an energetic barrier and ‘switch’ that holds SiaP in the closed conformation upon Neu5Ac binding (27). However, because the kink features in both our closed bound and mostly closed unbound structures, it may not impose a barrier to intrinsic closure, although it is straightened completely in structures generated by molecular dynamics (MD) simulations (discussed in detail later).

The prime candidate for signalling the hinge domains to open or close is the highly conserved triad of interacting residues R127, E186, and H209 (28), although this has been questioned in the literature (36). In the unbound state, R127 within the hinge region is proposed to form a strong interaction with E186, which is significantly weakened upon Neu5Ac binding and coordination by R127. This results in a stronger interaction between E186 and H209, which is also housed in the β-strand hinge area. This network has been implicated in triggering the observed conformational change, and H209 has also been implicated in the protein folding process (27). From a structural superposition of our crystal structures in different conformations, there does not appear to be any major changes in this network to support the proposed role in triggering the conformational change. However, in fully open structures obtained from relaxed MD simulations (next section) we observe that the interaction between E186 and H209 is weakened—the distance between the oxygen (OE2) of E186 and the nitrogen (NE2) of H209 increases from 2.7 Å to 4.0 Å. Together with our mostly closed structure with acetate bound at R127, we suggest that this residue interaction network is partly involved in stabilising the open and closed conformations of SiaP.

Superposition of the structures also reveals that the largest movements upon closure occur at surface residues near the substrate-binding site, particularly R50, N150, F170, and the residues that they interact with. In the closed-bound structure, R50 and N150 are conspicuously stacked together at the protein surface, forming one of the few new residue-residue contacts between the N- and C-terminal domains upon closure (as analysed by DynDom). We collectively refer to this entire region as a latch, which was also previously identified in the *H. influenzae* SiaP (45) and in the *P. profundum* SiaP (17). R50 undergoes the largest movement between the mostly closed-unbound and closed-bound states (~5 Å). Indeed, mutation of the equivalent latch residues (R49 and N148) in *Pp*SiaP showed a dramatic reduction in transport activity using the full SiaP-SiaQM system (17). The neighbouring residue D49 is also of interest here, as it coordinates both the glycerol moiety of Neu5Ac and R70. This, in turn, holds R70 in close contact with N150 across the binding cleft, with the guanidinium group of R70 hydrogen bonding to the N150 backbone carbonyl. Of the new residue-residue contacts upon closure, the R70-N150 contact is one of two H-bonding interactions formed between domains, the other being Q74-T155. Both interactions are observed in the mostly closed unbound and the liganded structures, however, in the mostly closed structure, the aforementioned R50-N150 stacking interaction is not present. Therefore, it appears that D49 is also involved in ligand sensing and protein closure, by stabilising the interaction between R70 (domain I) and N150 (domain II).

Examination of the Neu5Ac bound structure (**Figure 4C**) explains why *Aa*SiaP prefers Neu5Ac (*K*_D_ = 430 nM) over Neu5Gc (*K*_D_ = 4,400 nM). Focusing on the acetyl moiety of Neu5Ac (to which the hydroxyl of Neu5Gc is bound, **Figure 2C**) shows that it is closely coordinated by the backbone of A66 and F65, the sidechain of N214, and a conserved water that is part of the water network (**Supplementary Figure 6**). Neu5Gc binding would either require realignment of the molecule to accommodate the additional hydroxyl, potentially binding in a suboptimal conformation, or disruption of the water network, which is known to be important for binding.

In summary, the conformational changes of SiaP appear to involve: 1) direct and indirect (water-mediated) protein-ligand interactions, 2) the triad of residues connected to the β-strand hinge region, 3) an energetic barrier imposed by the kinked α9 helix, and 4) latching interactions between the two domains.

### MD simulations of the AaSiaP suggest an open state is favoured and correlates with hinge straightening

Given the possibility that crystal packing may affect the conformation of the protein, we employed full atom MD simulations over 500 ns to determine whether *Aa*SiaP sampled different conformations *in silico*. Our simulations started from three apo states of *Aa*SiaP to assess the conformational flexibility of the protein: open, mostly closed and fully-closed (with Neu5Ac removed); the fully-closed Neu5Ac-bound state, and the fully-closed Neu5Ac and conserved water-bound state. The degree of opening was assessed by measuring the distance between the Cα atoms of the aforementioned latch residues, R50 and N150 (**Figure 5A,B**).

**Figure 5.**
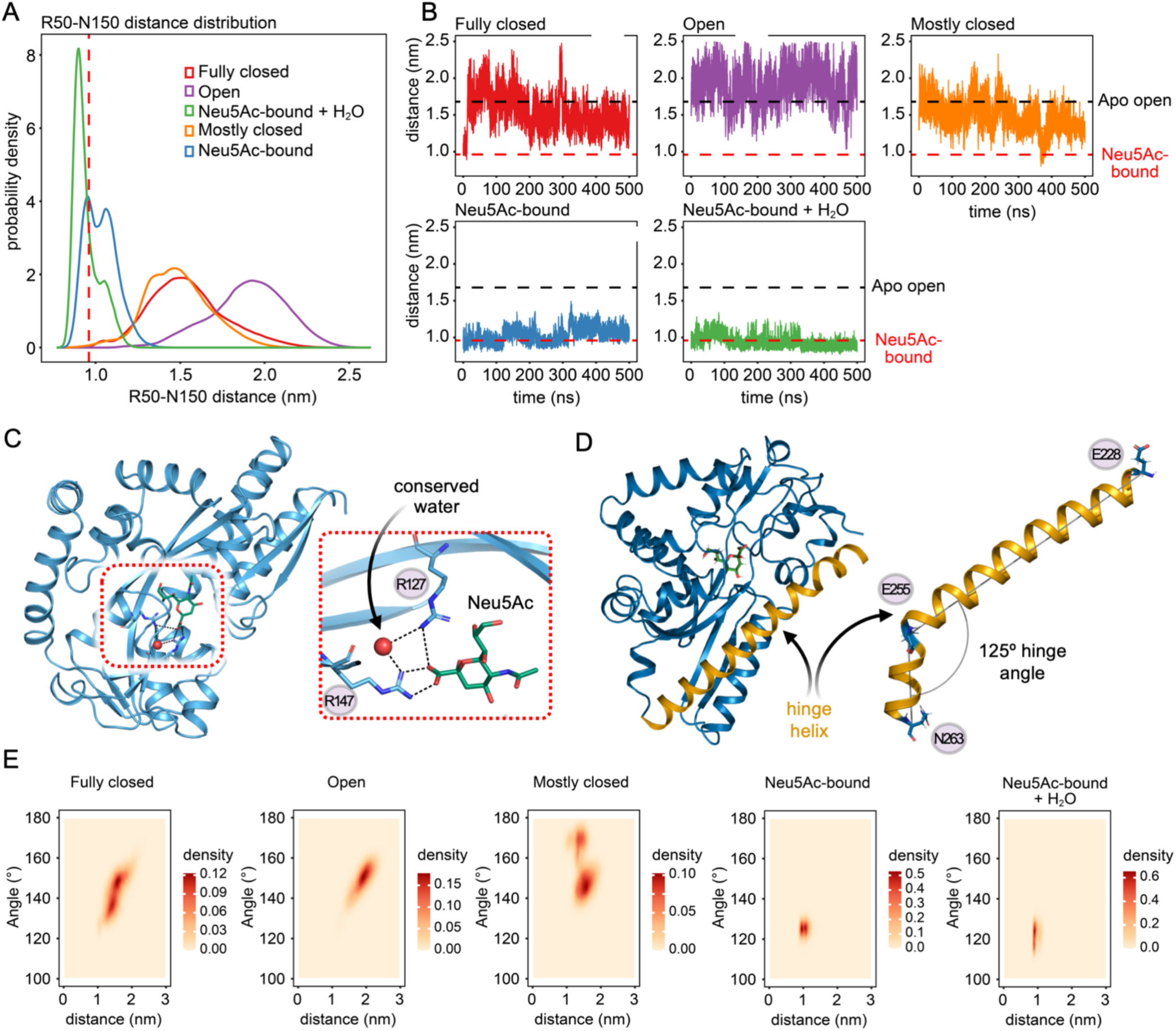
Molecular dynamic simulation results of *Aa*SiaP. **A**) The distribution of the distance between Cα atoms of R50 and N150, calculated from five independent 500 ns MD simulations. Simulations initiated from different conformations of *Aa*SiaP are shown in different colours: fully-closed (red), Neu5Ac-bound (blue), Neu5Ac-bound with crystallised waters (green), open (purple) and mostly closed (orange). The vertical dashed line indicates the corresponding Cα atom distance observed in the crystal structure of Neu5Ac-bound *Aa*SiaP. **B**) Dynamic changes in the distance between Cα atoms of R50 and N150 across the five simulations. The horizontal black and red dashed line represent the distances observed in the apo-open and Neu5Ac-bound crystal structures, respectively. **C**) Representation of the highly conserved water molecule within the binding pocket of the *Aa*SiaP Neu5Ac-bound conformation. The inset on the right is a cross-eyed stereo plot of the binding site showing the interaction of the water molecule with the twin arginine residues (R147 and R127) and the Neu5Ac. **D**) Representation of the hinge helix in the crystal structure of Neu5Ac-bound *Aa*SiaP. The inset on the right demonstrates the kink angle of the hinge helix, which is defined by the Cα atoms of E226, E255 and N263. **E**) The probability densities of distances between R50 and N150, alongside the kink angle (as shown in **D**).

The simulations starting from the fully-closed and mostly closed states sampled the open state with similar R50-N150 distance distributions (**Figure 5AB**, orange and red lines), whereas the simulation started from the open state sampled a broader distribution of much larger distances (**Figure 5AB**, purple line). Only the apo state simulation initiated from the mostly closed state spontaneously sampled closed state distances, with the simulation initiated from the fully-closed state never returning to closed-state distances, suggesting that while sampling of the closed state by the apo state of *Aa*SiaP in solution is possible, it is uncommon. In contrast, in the Neu5Ac-bound state simulations, the sampled R50-N150 distances are generally close to that of the Neu5Ac-bound crystal structure (**Figure 5AB**, blue and green lines). The second peak in the distance distribution for the simulation without crystallised water molecules in the binding site (blue line) corresponds to the later stages of the simulation where the binding pocket opens slightly, although the Neu5Ac remains bound. A highly conserved water molecule (*i.e.*, observed in other SiaP crystal structures) was found between the twin arginine residues of the binding site (R147 and R127) and the carboxyl oxygen of Neu5Ac (**Figure 5C**); the R50-N150 distances in the simulation that included this crystallographic water molecule more often correspond to the bound state (**Figure 5A,B**), suggesting that it may contribute to the affinity of Neu5Ac binding and thus the stabilisation of the fully-closed conformation. This is consistent with our crystallographic results above. We note that the water molecule in question is not a part of the Neu5Ac bound water shell (**Supplementary Figure 5**), but rather a structural water molecule. Previous studies demonstrate that both *Hi*SiaP-R147K (PDB id: 2xwi) and *Hi*SiaP-R147A (PDB id: 2xwk) weakly bind Neu5Ac *in vitro*, but in their crystal structures sialic acid is bound as in the wildtype and the protein is in the fully closed state. This indicates that the carboxylate-arginine interactions are not essential for closure, though the simulation data here show that the interaction helps to maintain or stabilise the fully closed state.

To determine whether the opening and closing motion of *Aa*SiaP is related to bending of the hinge helix α9, the probability densities of the R50-N150 distance and the helix kink angle (**Figure 5D**) were computed (**Figure 5E**). The narrow distance distribution observed in the Neu5Ac-bound simulations correlates with limited flexibility of the hinge helix, which predominantly adopted an angle of ~125°, even when the binding pocket opened in the simulation without crystallographic water molecules. In contrast, the ~125° angle of the hinge was only observed at the start of the ligand-free simulation initiated from the fully-closed conformation; otherwise, the apo state favoured larger hinge angles. Moreover, this transition to wider hinge angles and larger R50-N150 distances at the start of the apo-state simulation initiated from the fully-closed state was rapid, and while distances close to that measured for the closed Neu5Ac-bound state are sometimes sampled during the remainder of the simulation, ~125° hinge angles are not, implying an energy cost associated with maintaining this bound-like hinge angle. The apo-state simulations initiated from the fully-closed state had two favoured hinge angles of ~135° and ~150°, with the latter also favoured by the simulation initiated from the open state. In contrast, the simulation initiated from the mostly closed state favoured hinge angles of either ~145° or ~170°. Together, these results suggest that while hinge straightening is correlated with opening of the binding pocket, it is not a straightforward relationship, as the mostly closed state simulation sometimes samples larger hinge angles than the open state simulation, but the open state simulation generally samples longer R50-N150 distances, representing a more open binding pocket.

### Apo-AaSiaP is more open in solution than in the crystal structure

We used small angle X-ray scattering (SAXS) to investigate the conformational dynamics of *Aa*SiaP in solution, complementing the crystallographic data and molecular dynamic simulations. Although only a low-resolution technique (~10 Å resolution), SAXS is a label-free approach that is sensitive to small changes in hydrodynamic parameters such as the conformational change expected to occur upon ligand binding [method review here (46)].

We measured scattering data collected in the presence and absence of Neu5Ac. Addition of Neu5Ac resulted in a reduction in the radius of gyration (*R*_g_) and the maximum interatomic distance (*D*_max_): without Neu5Ac the *R*_g_ = 21.31 ± 0.06 Å and the *D*_max_ = 68.1 Å, whereas in with 1 mM Neu5Ac the *R*_g_ = 20.43 ± 0.07 Å and the *D*_max_ = 65.0 Å, and with 10 mM Neu5Ac the *R*_g_ = 20.08 ± 0.02 Å and the *D*_max_ = 60.7 Å (**Table 3**). The significant decrease observed in *R*_g_ and *D*_max_ values from an absence of Neu5Ac in solution to saturating Neu5Ac in solution supports that the protein undergoes a significant detectable conformational change upon ligand binding.

**Table 3.** Summary of structural parameters for *Aa*SiaP from small angle X-ray scattering experiments.

| SAXS data collection parameters |  |  |  |  |  |
| --- | --- | --- | --- | --- | --- |
| instrument | Australian Synchrotron SAXS/WAXS beamline |  |  |  |  |
| detector | PILATUS 1M (Dectris) |  |  |  |  |
| wavelength (Å) | 1.0332 |  |  |  |  |
| maximum flux at sample | $8 \times 10^{12}$ photons per second at 12 keV | | | | |
| camera length (mm) | 1600 |  |  |  |  |
| q-range (Å<sup>-1</sup>) | 0.006–0.5 |  |  |  |  |
| exposure time | Continuous 1-s frame measurements |  |  |  |  |
| sample configuration | SEC-SAXS with co-flow |  |  |  |  |
| sample temperature (°C) | 12 |  |  |  |  |
| SAXS data analysis | Apo | Neu5Ac (0.1 mM) |  | Neu5Ac (10 mM) |  |
| *Guinier analysis* |  |  |  |  |  |
| $I_o$ (cm<sup>-1</sup>) | $0.029 \pm 4.5 \times 10^{-5}$ | $0.029 \pm 5.7 \times 10^{-5}$ | | $0.053 \pm 3.1 \times 10^{-5}$ | |
| $R_g$ (Å) | $21.31 \pm 0.06$ | $20.43 \pm 0.07$ | | $20.08 \pm 0.02$ | |
| *P(r) analysis* |  |  |  |  |  |
| $I_o$ (cm<sup>-1</sup>) | 0.03 | 0.03 | | 0.05 | |
| $R_g$ (Å) | 21.30 | 20.33 | | 19.99 | |
| $D_{max}$ (Å) | 68.1 | 65.0 | | 60.7 | |
| Porod volume (Å<sup>-3</sup>) | 47313.3 | 51858.8 |  | 51187.3 |  |
| *CRY SOL analysis* |  |  |  |  |  |
| open (X<sup>2</sup> value) | 0.708/0.949 | 0.497/0.513 |  | 7.317/11.067 |  |
| mostly closed (X<sup>2</sup> value) | 2.203/2.291 | 0.759/0.775 |  | 0.375/0.482 |  |
| closed (X<sup>2</sup> value) | 2.583 | 0.873 |  | 0.366 |  |
| *SAXS-MoW2<sup>a</sup>* |  |  |  |  |  |
| estimated MW (kDa) | 28.5 | 23.2 |  | 28.1 |  |
| Crysol prediction | chain C<br>(open) | chain D<br>(open) | chain A<br>(closed) | chain B<br>(closed) | bound *Aa*SiaP |
| $R_g$ (Å) | 20.48 | 20.22 | 20.07 | 19.99 | 19.7 |
| $D_{max}$ (Å) | 72.17 | 70.93 | 71.13 | 68.14 | 66.81 |

<sup>a</sup>http://saxs.ifsc.usp.br/

The experimental scattering data were fit with theoretical scattering curves calculated from the X-ray crystal structures (**Figure 6A,B**). As expected, the data collected with Neu5Ac (10 mM) were best fit by the closed structure, whilst data without Neu5Ac were initially best fit by the open structure. Fitting fully open MD-based models (presented in the previous section) to our SAXS data gave an even better fit than the open structure (**Figure 6B**). Further analysis of the fully open MD model corroborates our previous hypothesis that R50 and N150 form part of the latch, which moves significantly further apart from closed, to mostly closed, to open, to fully open structures (**Figure 6C**). Overall, our SAXS and MD analyses support that the ensemble average of *Aa*SiaP conformations sampled in solution is more open than the open conformation determined in the crystal structure, although the MD analysis suggests that the open conformation can (rarely) sample the closed conformation.

**Figure 6.**
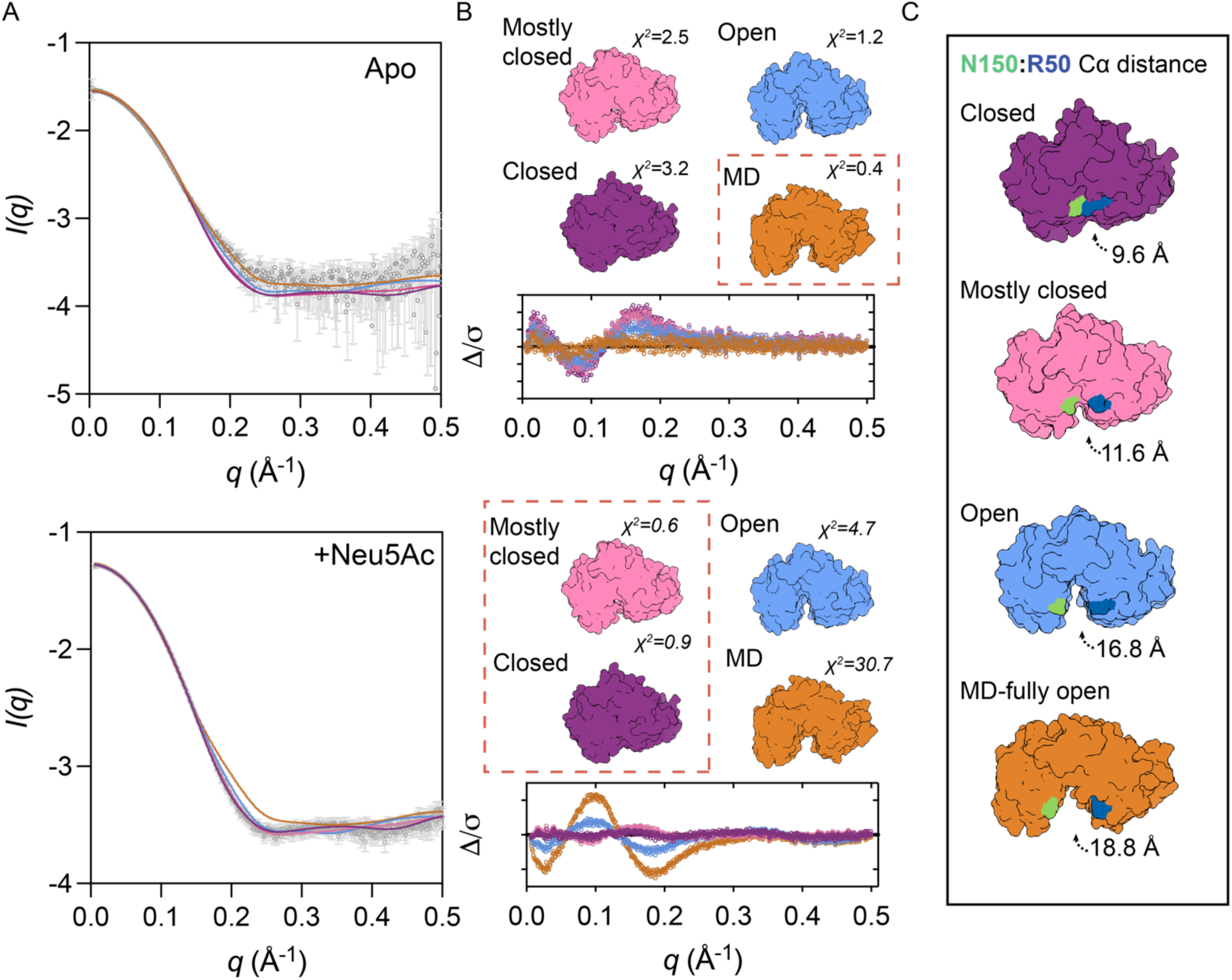
Small angle X-ray scattering analysis of *Aa*SiaP. **A**) *Aa*SiaP scattering data (grey), fitted with the theoretical scattering curves from various models using *Crysol* (70). Data collected in the absence of Neu5Ac (top, ligand-free); data collected in the presence of 10 mM Neu5Ac (bottom). **B**) Error-weighted residuals plots of the fits, alongside the structural models used to generate the fits. The best fits to the data (lowest χ^2^) are indicated by the red dashed box. **C**) Illustrating the large extent of the conformational change between the closed structure and a fully open MD model. The distance displayed is the distance between the C_α_-atoms of R50 and N150, a residue pair proposed to latch together (45).

**Figure 7.**
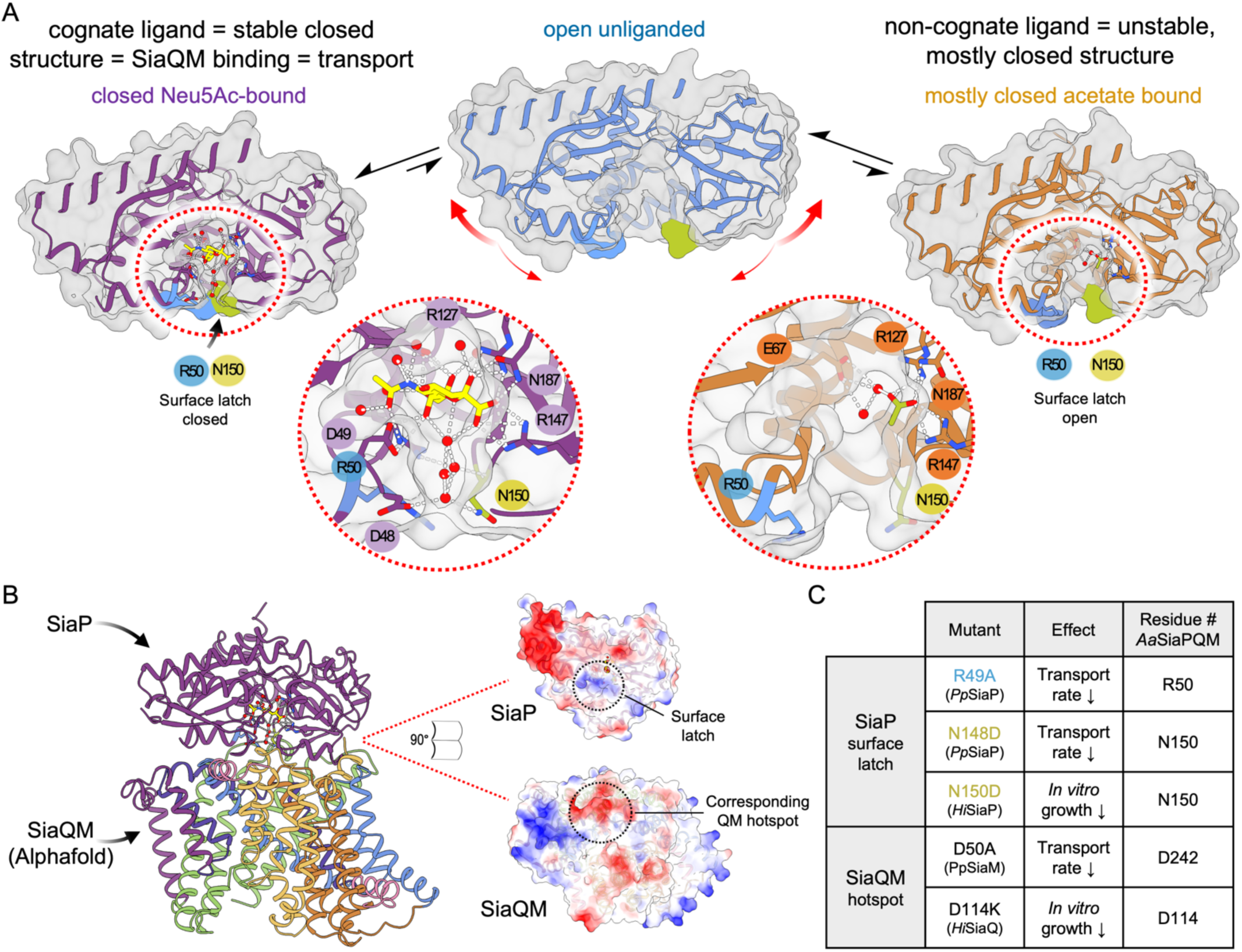
Full closure of SiaP involves latching and a water network that extends to the interaction surface. **A**) The open unliganded form of *Aa*SiaP can sample a range of conformations including (largely) a more open structure that found in the crystal, and (rarely) the closed conformation. Binding Neu5Ac and a water network stabilises the closed conformation that can bind to the SiaQM membrane components. In contrast, binding a non-cognate ligand leads to a mostly closed conformation, without a stabilised water network and presumably cannot bind the SiaQM membrane components. **B**) Alphafold2 modelling of SiaP-SiaQM, with an open-book representation of the interacting surfaces. **C)** Table showing the functional effects of mutating residues at either the surface latch region, or the corresponding region on SiaQM.

## Discussion

### New structures suggest small molecules can induce only partial closure of AaSiaP

We report two crystal structures of *Aa*SiaP that show three conformations: 1) a fully closed conformer with Neu5Ac bound, 2) an open conformer that is similar to previously reported apo structures, 3) and a unique apo conformer that is mostly closed. The apo-*Aa*SiaP crystal structure contains two conformations of *Aa*SiaP in an uncommon space group (C2) amongst SiaP structures, shared by only one other structure deposited in the PDB with four unique conformations, where one of four monomers in the asymmetric unit is bound with the sialic acid analogue Neu5Ac2en [PDB ID: 2CEX, (27)]. Apo-*Aa*SiaP crystallised in acetate-containing conditions (200 mM) and the electron density in the binding site is more consistent with acetate than a water molecule or Neu5Ac. The crystal contacts/interfaces formed in the apo-*Aa*SiaP structure are largely the same and therefore unlikely to have induced the observed difference in conformation.

We propose that the binding of acetate can stabilise only partial closure, which is consistent the induced fit model of substrate binding in that the non-cognate ligand is unable to induce the tight binding (*i.e.*, closed) conformation. The specificity and sensitivity of SiaP for Neu5Ac has been well-established in the literature, with even small modifications to the Neu5Ac structure having significant effects on binding affinity (27). Closure around a non-cognate ligand is consistent with models proposed for other substrate-binding proteins, where their binding induces a partially closed conformation that is ‘conformationally mismatched’ to the membrane domains, so does not facilitate transport (47). Recently, Peter et al. (2021) reported a structure of *Vc*SiaP with a peptide bound, further supporting that SiaP can bind non-cognate ligands. The authors sensibly propose that full closure is triggered by the physical bridging of the two domains by the ligand. Our mostly closed *Aa*SiaP structure with acetate bound suggests that the carboxylate interaction may be sufficient to induce closure, but not a full ‘latching’ between domains.

Considering now the proposed mechanism of TRAP transporter import, we suggest that failure to induce full closure by non-cognate ligands is a method of discrimination for SiaPQM systems, in that only the cognate ligand induces a fully closed conformation that can productively interact with the membrane domains to facilitate transport, which has been proposed in the ABC family transporters (47).

### Apo-AaSiaP displays conformational heterogeneity and closure in solution

Our MD simulations demonstrate that the apo state strongly favours an open state—in terms of both the R50-N150 distance and the α9 helix hinge angle—regardless of the structure from which the simulation is initiated. However, the R50-N150 distances are often larger than in the apo open-state crystal structure and that near closed conformations can be sampled, although rarely. The small angle X-ray scattering data also suggest that in solution at biologically relevant temperatures, ligand-free *Aa*SiaP exists predominantly in an open conformation that is likely more open than in the open-state crystal structure. This is consistent with findings from PELDOR spectroscopy experiments on frozen *Vc*SiaP (36) and atomistic MD simulations based on this PELDOR data, which suggest that *Vc*SiaP in solution may also be more open than in the open-state crystal structure (48). More recently, rare closure events have been reported for an apo disulfide engineered *Hi*SiaP construct, where residues within the latch region of SiaP are substituted (S15C/A194C) (49). In this experiment, disulfide formation between these residues only occurs when SiaP closes, and the residues were in close contact in the required geometry and was detected as a distinct band shift by non-reducing SDS-PAGE in the absence of oxidising agents. A time-course experiment based on this band shift and quantifying the relative proportions of non-sulfide and disulfide-linked proteins suggests that the apo *HiS*iaP protein underwent an intrinsic closing motion approximately every 23 seconds. This is equivalent to less than 0.01% of all molecules closing within one millisecond, and since the apo SiaP quickly reopens in solution, only a small fraction of closed apo SiaP are present at one time. This is generally supported by our MD simulations, which only show partial closure over the 500 ns simulation time.

In contrast, the closed conformation is strongly favoured when Neu5Ac is bound, but the conserved water molecule between R127 and R147 is required to fully stabilise this state. This data fits nicely with recent work that demonstrates the importance of water networks for Neu5Ac binding (25). Our crystal structure of the Neu5Ac bound shows a highly ordered network of waters in the ligand binding site that coordinate both the ligand and the protein. Many of these waters highly conserved when compared to other Neu5Ac bound SiaP structures.

### Implications for the formation of the SiaP-SiaQM complex and substrate release

A prominent model proposed by Mulligan et al. (2009) suggests that only the closed-bound SiaP conformation productively binds the SiaQM membrane domains, which in turn modulate SiaP into an open conformation, prompting transfer of the substrate (24). More recently, Peter et al. (2021) reported closure and binding of SiaP to peptides based on the predicted periplasmic loops of the SiaQ subunit. These findings are consistent with the ‘scoop-loop model’ of some ABC transporters, where the periplasmic loops of the membrane domains help dislodge the bound substrate and/or prevent the released substrate reassociating with the substrate-binding protein instead of the membrane domains (50). However, modelling of the SiaP-SiaQM interaction does not point to an obvious scoop-loop, even considering large elevator-type movements of the transport domain (17). We speculate that it is more likely that substrate release is governed by SiaP-SiaQM surface interactions that disrupt the Neu5Ac binding site allosterically.

Evidence that the SiaP and SiaQM domains are conformationally coupled has come from Peter et al. (2024), who assessed interactions for *Hi*SiaQM disulfide engineered constructs locked in the inward-facing state (IFS) and outwards-facing state (OFS) mixed with both wildtype and the disulfide engineered *Hi*SiaP constructs (described above). Their findings largely support a transport model where the closed SiaP preferentially interacts with the SiaQM IFS and an apo open SiaP preferentially interacts with the SiaQM OFS. Exactly how SiaP interacts with the OFS needs investigation, and in particular whether or not SiaP is fully open during this interaction. It may be possible that SiaP adopts an intermediate open state while bound to the OFS, considering that the closed SiaP could still occasionally interact with the OFS (49). How the fully open state observed here fits in with the mechanism is unclear, though we speculate that this state should have the lowest affinity for SiaQM, and that this promotes dissociation from the transporter. Further structural and functional studies are needed to better assess these interactions.

Our data suggests conserved water networks and residues involved in co-ordinating Neu5Ac are required to give the fully closed P domain, which presumably has the highest affinity for the inward-facing SiaQM domains. We report partial closure with co-ordination of the non-cognate ligand acetate to key residues within the Neu5Ac binding site, but this partial closure was insufficient to induce latch closure. Residues involved in this latch are critical to interactions with the SiaQM domains and for transport activity (17). Overall, our data support that the substrate-binding induced latch formation may be a key requisite of high affinity SiaP docking and subsequent substrate handover to SiaQM.

To conclude, we report a mostly closed-non-cognate bound conformation of *Aa*SiaP similar to structures previously reported for ABC substrate-binding proteins (32, 47, 51), but which has not to our knowledge been observed for TRAP substrate-binding proteins. We propose that interactions between residues at the resulting domain interface stabilise the mostly closed-unbound conformation in equilibrium, and that ligand-binding induces small but important residue movements that mediate productive interactions with the membrane domains. These results are inconsistent with a simple two-state induced-fit model of ligand binding; thus, we instead propose a special case of induced fit that involves an intrinsic conformational change in the absence of ligand. Our findings contribute to a deeper understanding of the conformational dynamics of SiaP proteins and other TRAP substrate-binding proteins, which may aid in the design of novel inhibitors for use as antimicrobial drugs.

## Experimental procedures

### Multiple sequence alignment

The position-specific iterated protein (PSI-)BLAST was performed using the *A. actinomycetemcomitans* SiaP protein sequence (NCBI accession # WP_005538762.1) as the query to identify homologous substrate-binding proteins. Twelve of the highest-scoring sequence identity sequences from a diverse range of species, including four proteins with available PDB structures (**Figure 1**), were chosen for sequence alignment and structural comparison (**Supplementary Figure 1**): NCBI accession no. WP_109077085.1, 2CEY_A, WP_077422743.1, WP_100297742.1, WP_132021368.1, WP_039082755.1, 4MMP_A, WP_075321769.1, WP_048716221.1, WP_103853354.1, 4MNP_A, 4MAG_A. Multiple sequence alignment was performed using Clustal Omega (https://www.ebi.ac.uk/Tools/msa/clustalo/) (52) and the figure was produced in ESPript 3.0 (http://espript.ibcp.fr/ESPript/cgi-bin/ESPript.cgi) (53).

### Transformation and protein overexpression

The *A. actinomycetemcomitans* strain 624 *siaP* gene (NCBI accession no. NZ_CP012959, Region: 361430–362416) was synthesised by Genscript and ligated into expression vector pET22B(+). In this construct the native periplasmic signal peptide sequence, as predicted by the SignalP-5.0 server (54), was replaced with *pelB* periplasmic signal peptide sequence from *Erwinia carotovora*. A residual two-residue (methionine and aspartate) cloning fragment was retained at the N-terminus of the mature chain following removal of the 22-residue *pelB* leader sequence. Chemically-competent *E. coli* BL21 (DE3) cells were then transformed with recombinant pET22B(+) containing the *A. actinomycetemcomitans siaP* sequence and allowed to recover in super optimal broth with catabolite repression at 37 °C for 1 h. Recombinant cells were then plated on Luria-Bertani (LB) agar medium supplemented with ampicillin (100 μg/mL) and incubated overnight at 37 °C. After being confirmed, the recombinant strain was stored at −80 °C.

Starter cultures of the recombinant strain were prepared from a single colony, which was inoculated into LB medium supplemented with ampicillin (100 μg/mL) and incubated for 4 h at 37 °C and 180 rpm. Following incubation, the culture was centrifuged at 4,230 × *g* for 10 min and the resulting cell pellet washed with 1 × M9 salts, then inoculated into M9 medium supplemented with ampicillin (100 μg/mL) and incubated overnight at 37 °C and 180 rpm. For protein expression assays, a starter culture was used to inoculate a 4 L volume of M9 minimal medium supplemented with ampicillin (100 μg/mL), which was then incubated at 26 °C for ~3 h to an optical density at 600 nm of 0.3. To induce protein expression, isopropyl β-d-1-thiogalactopyranoside was added to a final concentration of 1 mM, and the culture incubated at 16 °C overnight. Following incubation, the cells were collected by centrifugation at 5,600 x *g* for 10 min and resuspended in periplasmic extraction buffer (50 mM Tris-HCl, 500 mM sucrose, 1 mg/mL lysozyme, pH 8.0) at a concentration of 5 mL/g cell weight. DNase (1 µg/mL) was added to the cell suspension and the mixture was incubated for 1 h at 4 °C. Following incubation, 10 mL of ice-cold water was added, and the lysate further incubated for 30 mins.

### Protein purification

All purification steps were performed at 4 °C, and samples were maintained on ice throughout. Anion-exchange chromatography (AEX) was performed using a Q Sepharose Fast Flow Column (HiPrep Q FF 16/10; GE Healthcare) pre-washed with three column volumes of AEX Buffer B (20 mM Tris-HCl, 1 M NaCl, pH 8.0) to elute bound proteins. The column was then equilibrated with three column volumes of AEX Buffer A (20 mM Tris, pH 8.0). The collected periplasmic fraction was loaded onto the column and bound recombinant *Aa*SiaP was eluted using an increasing gradient of AEX Buffer B over 10 column volumes.

Hydrophobic interaction chromatography (HIC) was then conducted using a Phenyl Sepharose 6 Fast Flow Column (HiPrep Phenyl FF (High Sub) 16/10, GE Healthcare) pre-washed with three column volumes of HIC Buffer B-SEC Buffer (20 mM Tris-HCl, 150 mM NaCl, pH 8.0) and equilibrated with three column volumes of HIC Buffer A (20 mM Tris, 1 M (NH_4_)_2_SO_4_). Ammonium sulphate (final concentration, 1 M) was added to the pooled *Aa*SiaP-containing fractions, which were then loaded onto the Phenyl Sepharose column. Bound *Aa*SiaP was again eluted using an increasing gradient of HIC Buffer B over 10 column volumes.

Size-exclusion chromatography (SEC) was employed as the final purification step using a Superdex 200 Column (Superdex 200 Increase 10/300 GL, GE Healthcare) pre-equilibrated with three column volumes of SEC buffer (Tris-NaCl). The pooled fraction from the HIC step was concentrated to ~40 mg/mL (Millipore) then loaded onto the column. *Aa*SiaP-containing fractions were pooled, and purification was confirmed by sodium dodecyl sulphate polyacrylamide gel electrophoresis analysis. Purified protein that was not used immediately was flash-cooled in liquid nitrogen and stored at −80 °C.

### Differential scanning fluorimetry

Purified *Aa*SiaP in SEC buffer was concentrated using Vivaspin centrifugal concentrators (10,000 Da molecular weight cut-off; Sartorius) at 8000 × *g* then subjected to two rounds of buffer exchange using equimolar Tris-KCl buffer and a 5-mL desalting column (HiTrap Desalting; GE Healthcare) pre-equilibrated with four column volumes of Tris-KCl. Differential scanning fluorimetry was conducted as described (55), with some modifications, using a QuantStudio 3 Applied Biosystems real-time polymerase chain reaction (PCR) system with differential scanning fluorimetry capability (Thermo Fisher Scientific). Assays were repeated in triplicate in a 96-well 0.2-mL PCR-compatible plates. The 20 μL reaction volumes contained 16 μL of Tris-NaCl/KCl buffer (+/− Neu5Ac at various concentrations), 1 μM *Aa*SiaP (2 μL), and 50 × SYPRO orange dye (2 μL). Neu5Ac was dissolved in small aliquots of buffer in triplicate to achieve concentrations ranging from 0.00025–1 mM, corresponding to the expected *K*_D_ range. Samples were gradually cooled from an initial temperature of 29 °C to 20 °C and held for 1 min. Samples were then heated incrementally to 95 °C at a rate of 0.015 °C/s (~84 mins), and fluorescence intensity was recorded over time.

### Fluorescent labelling and microscale thermophoresis

For subsequent microscale thermophoresis (MST) binding experiments, *Aa*SiaP was labelled at lower pH, which directed fluorescein 5-isothiocyanate (FITC) towards the lower pKa alpha amine at the N-terminal. This was done to minimise the amount of structural and conformational perturbation, as the protein contains 29 lysine residues that would be more reactive at higher pH. *Aa*SiaP (30 µM) in storage buffer (Tris-HCl 50 mM, NaCl 150 mM, pH 8) was buffer exchanged into PBS, pH 7.4, incubated with FITC (1:40 molar ratio *Aa*SiaP:FITC in DMSO) for two hours at room temperature, then spin concentrated iteratively (3kDa molecular weight cut-off) to remove free dye before separation of the *Aa*SiaP-FITC conjugate by SEC, as detected by absorbance at 280 nm and 490 nm. Fluorophore/protein (F/P) molar ratio was determined from absorption at these wavelengths and calculated by the equation: Molar F/P = (A_495_ x C)/(A_280_ – [(0.35 x A_495_)], where C is a constant for the protein: C = (MW x E0.1%)/(389 x 195) = 34,000 g/mol x 0.693/389 x 195 = 0.3106), which gave a F/P ratio of 0.42. MST binding experiments were performed on a Monolith NT.115 (NanoTemper Technologies) with Monolith NT.115 Capillaries. Binding checks were performed with various concentrations of *Aa*SiaP with both Neu5Ac and Neu5Gc, which suggested that 10 mM Neu5Ac (100-fold higher than the highest concentration in the binding assay) suggested ligand autofluorescence at this excess concentration; low fluorescent counts were also observed for some replicates at lower concentrations, which we attribute to the low labelling ratio, so a higher concentration of 75 nM *Aa*SiaP was selected for binding affinity experiments. Binding affinity experiments were performed in duplicate (Neu5Gc) or triplicate (Neu5Ac) at 16 different concentrations of ligand, ranging from 100 µM to 0.00305 µM, and analysed with MO Affinity Analysis V2.3 software.

### Analytical ultracentrifugation

Sedimentation velocity analytical ultracentrifugation was performed using a Beckman Coulter ProteomeLab XL-1 Protein Characterisation System, with a Beckman Coulter AN50Ti rotor with sapphire and quartz windowed cells. Data were collected at 20 °C with absorbance optics (280 nm) and run at 42,000 rpm. Recombinant *Aa*SiaP at 0.42 mg/mL (12 μM, concentration measured by absorbance at 280 nm) was analysed in Tris-NaCl buffer in the absence of Neu5Ac, and with two concentrations of Neu5Ac (0.1 mM and 0.5 mM).

Data were analysed using UltraScan 4.0, release 2578 (56). Sedimentation data were evaluated by the two-dimensional spectrum analysis (2DSA) (57, 58), with simultaneous removal of time- and radially invariant noise contributions and fitting of boundary conditions. In addition, 50 Monte-Carlo iterations were performed on the 2DSA data set and 2DSA-Monte-Carlo solutions were subjected to parsimonious regularisation by genetic algorithm analysis (59). All fitting procedures were completed using the UltraScan LIMS cluster. Each-genetic algorithm model was visually assessed to ensure a good fit and a low RMSD using the FE Model Viewer utility in UltraScan (**Supplementary Figure 4**). Predicted values for hydrodynamic parameters were estimated in *Hullrad* (60) for *Aa*SiaP crystal structures.

### Crystallisation and structure determination

Crystallisation trials were set up using the Clear Strategy, ShotGun (SG1), and PACT premier screens in 96-well plates. Drops consisting of 400 nL of mother-liquor and protein solution (*Aa*SiaP at 20 mg/mL along with and without 0.75 mM Neu5Ac in SEC buffer) were mixed using the Mosquito Protein Crystallization System and the sitting-drop vapour-diffusion method and incubated at 20 °C. Apo *Aa*SiaP crystals that grew in SG1 conditions C11 (0.2 M sodium acetate trihydrate, 0.1 M sodium cacodylate, pH 6.5, 30% (w/v) PEG 8000) and F2 (0.2 M sodium acetate trihydrate, 0.1 M sodium HEPES, pH 7.5, 25% (w/v) PEG 3350), which were mounted in loops and preserved in 15% (v/v) cryoprotectant (50% (v/v) ethylene glycol, 50% (v/v) glycerol). Neu5Ac (0.75 mM) bound *Aa*SiaP crystals including grew in SG1 condition H2 (30% w/v PEG 4000) were not cryo-protected before being flash-cooled in liquid nitrogen prior to data collection.

The diffraction data for recombinant *Aa*SiaP were collected at the Australian Synchrotron (Melbourne, VA) on the MX2 beamline using an X-ray wavelength of 0.91840 Å equipped with an EIGER 16 M detector. The apo crystals from SG1 condition C11 diffracted to a maximum resolution of 2.58 Å and belonged to space group *C*2. In contrast crystals grown in SG1 F2 diffracted to a maximum resolution of 3 Å and also belonged to space group *C*2. Crystals from SG1 H2 diffracted to a maximum resolution of 1.90 Å and belong to space group *P*2_1_. Diffraction data were scaled and processed using *XDS* (60) and *Aimless* (*CCP4i2 suite*) (61). Resolution cut-offs were determined following the criteria that the CC_1/2_ of each dataset is above 0.35 and where the *I/σI* was equal or greater than 1.0. Data collection statistics are reported in **Table 2**.

Structures were determined by molecular replacement using *Phaser* (CCP4i2 suite) (63). Apo *Aa*SiaP was solved with four per asymmetric unit, two in an open conformation and two in a near-closed conformation, using two different search models: an apo-structure of *H. influenzae* SiaP (PDB ID: 2CEY) edited using *Sculptor* (CCP4i2 suite), and the structure of Neu5Ac-bound *Aa*SiaP. The structure of Neu5Ac-bound *Aa*SiaP was solved by molecular replacement using a liganded structure of *H. influenzae* SiaP (PDB ID: 2WYK) with two protein molecules in the asymmetric unit. For both structures, initial manual model rebuilding was done in *Coot* (64) and refined using *Refmac* (CCP4i2 suite) (65, 66). The resulting data was then exported to *Phenix* (67, 68) for refinement, with an initial round of refinement performed simulated annealing (cartesian) to reduce model bias. Refinement with translation-libration-screw (TLS groups were determined automatically using *Phenix*). Water molecules were identified and added in the later stages of refinement. The final model for apo-*AaSiaP* included two monomers in the open conformation (chains C and D) and two monomers in a near-closed conformation (chains A and B). Weak density was observed for cloning fragments at residues 1 and 2 at the N-terminus. Refinement statistics are presented in **Table 2**.

### Small angle X-ray scattering

Purified *Aa*SiaP in Tris-NaCl (SEC buffer) was spin-concentrated 8,000 × *g* using Vivaspin centrifugal concentrators (10,000 Da molecular weight cut-off; Sartorius) before being subjected to buffer exchange into SAXS buffer (20 mM Tris, 50 mM NaCl, pH 8) using a 5 mL HiTrap desalting column pre-equilibrated with three column volumes of SAXS buffer.

Small angle X-ray scattering (SAXS) data were collected on the SAXS/WAXS beamline equipped with a Pilatus 1M detector (170 × 170 mm, effective pixel size, 172 × 172 μm) at the Australian Synchrotron (69). A sample detector distance of 1600 mm was used, providing a *q* range of 0.05–0.5 Å^−1^. Here 70 µL of purified *Aa*SiaP protein at 10 mg mL^−1^ was injected onto an inline Superdex S200 Increase 5/150 GL (GE Healthcare) SEC column, equilibrated with 20 mM Tris–HCl pH 8.0, 150 mM NaCl, supplemented with 0.1% (*w*/*v*) sodium azide, using a flow rate of 0.45 mL min^−1^. For data collected in the presence of Neu5Ac, the same buffer was used with Neu5Ac added to a final concentration of 10 mM. Scattering data were collected in one second exposures (λ = 1.0332 Å) over a total of 400 frames, using a 1.5 mm glass capillary, at 8 °C. 2D intensity plots were radially averaged, normalised to sample transmission, and background subtracted using the *Scatterbrain* software package (Australian Synchrotron), and then analysed using *Chromixs* in the ATSAS suite (70). The theoretical scatter, *R_g_* and *D_max_* values for the open, near-closed, and closed-liganded conformations were calculated in *Crysol* (71) and compared to the experimental data.

Prior to *ab initio* shape determination, the ambiguity score for each condition was calculated in *Ambimeter* (72); in all cases the score was less than 1.5 (1.146, 0.9542, and 0.7782 for 0, 0.1, and 10 mM Neu5Ac respectively) indicating that unique non-ambiguous shapes could be determined from the experimental scatter. Ab initio bead models were then created in ten independent *Gasbor* (73) runs (accessed *via* https://www.embl-hamburg.de/biosaxs/atsas-online/) and the representative models, as determined by *Damaver* and *Damfilt*, were superimposed onto the *Aa*SiaP crystal structures using *Subcomb* (74) *via* the *SASpy* plugin (75) for open source *PyMOL*. The parameters for data collection and processing are summarised in **Table 3**.

### Molecular dynamics simulations

All simulations were performed using the GROMACS 2021.5 software package (76) and the CHARMM36m force field (77); the integration time-step was 2 fs, and periodic boundary conditions were applied. The parameters for Neu5Ac were obtained from the CHARMM36m ligand database. Short-range electrostatic and van der Waals interactions were cut-off at 1.2 nm; the particle mesh Ewald method (78) as used to treat long-range electrostatic interactions and the van der Waals interactions were force-switched from 1.0 to 1.2 nm. Non-bonded neighbour-lists were updated using the Verlet cutoff-scheme (79). The LINCS algorithm (80) was used to constrain the hydrogen bond lengths to their equilibrium value. The temperature was maintained at 298 K and the pressure at 1 bar using the velocity-rescale thermostat (81) and the Berendsen barostat (82), respectively, aside from during the reduction simulation, which used the Parrinello-Rahman barostat (83). Each production simulation lasted 500 ns, with coordinates saved every 5 ps.

#### Simulation preparation

In all systems, the energy was first minimised using the steepest descent algorithm until the maximum force changed by less than 1000 kJ·mol^−1^·nm^−1^, and then the system was solvated using the TIP3P water model (84) (**Supplementary Table 4**). Na^+^ and Cl^−^ ions were then added to neutralise the system (**Supplementary Table 4**), followed by again minimising the systems energy. Initial velocities of each system were randomly generated from a Maxwell-Boltzmann distribution at 50 K in the NVT ensemble, followed by a 10 ps equilibration and then heating smoothly from 50 K to 298 K over 210 ps, and a second brief equilibration at 298 K for 40 ps. Each system was then equilibrated for 250 ps with the temperature and pressure maintained at 298 K and 1 bar.

#### Analysis

Simulation trajectories were visualised using Visual Molecular Dynamics (85) and images made using Pymol (The PyMOL Molecular Graphics System, Version 3.0 Schrödinger, LLC.). Distances and angles were calculated using the GROMACS analysis tools *gmx mindist* and *gmx gangle*, respectively, and the probability densities calculated and plotted using *R 4.3.2* (86). The minimum distance was between the Cα atom of residues R50 and N150, and the helix kink angle was defined as the angle between the vectors connecting the Cα atoms of residues N263-E225 and E225-E228.

## Acknowledgements

This research was undertaken in part using the MX2 beamline at the Australian Synchrotron, part of ANSTO, and made use of the Australian Cancer Research Foundation (ACRF) detector as well as the SAXS beamline at the Australian Synchrotron, part of ANSTO. R.C.J.D. acknowledges the following for funding support, in part: 1) the Marsden Fund council from Government funding, managed by Royal Society Te Apārangi (contract UOC1506); 2) a Ministry of Business, Innovation and Employment Smart Ideas grant (contract UOCX1706); and 3) the Biomolecular Interactions Centre (UC). R.A.N. and J.RA. also acknowledges the Marsden Fund council from Government funding, managed by Royal Society Te Apārangi (contract UOC1506).

## Supporting information

This article contains supporting information.

## Contributions

**TRKH:** Conceptualisation, formal analysis, investigation, methodology, visualisation, writing— original draft, review & editing, finalising manuscript.

**JD:** Conceptualisation, formal analysis, investigation, methodology, project administration, resources, validation, supervision, writing—original draft, review & editing, finalising manuscript.

**SQ:** formal analysis, investigation, methodology.

**MC:** Conceptualisation, resources, validation, supervision, writing—original draft, review & editing.

**ZT:** formal analysis, investigation, methodology.

**JC:** formal analysis, investigation, methodology, writing—review & editing.

**SP:** formal analysis, investigation, methodology.

**RF:** Resources, writing—review & editing.

**JRA:** formal analysis, investigation, methodology, writing—review & editing.

**RN:** Conceptualisation, resources, writing—review & editing.

**RCJD:** Conceptualisation, funding acquisition, project administration, formal analysis, resources, supervision, review & editing.

## Supporting Information

### Supplementary Tables

**Supplementary Table 1.**
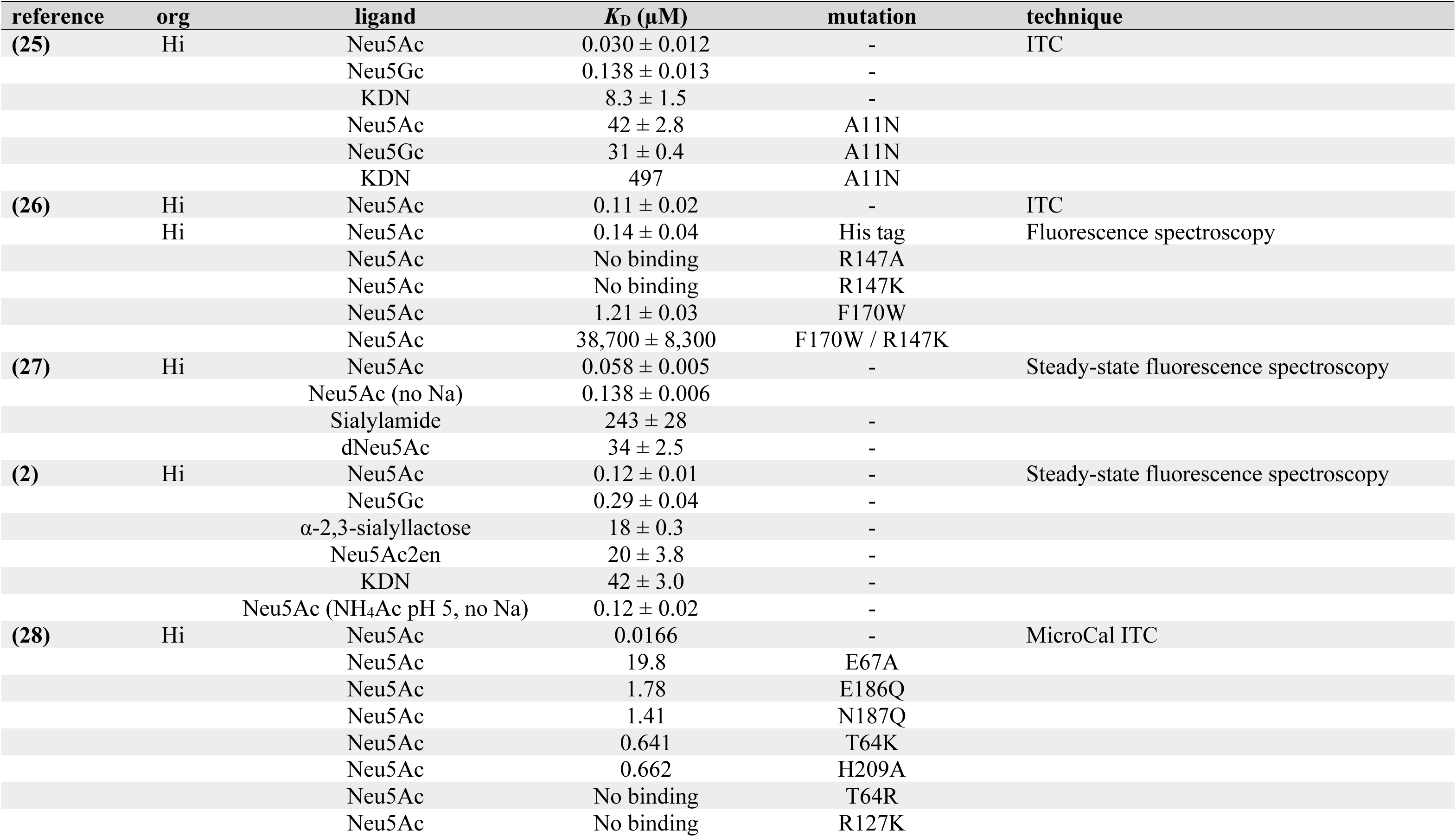
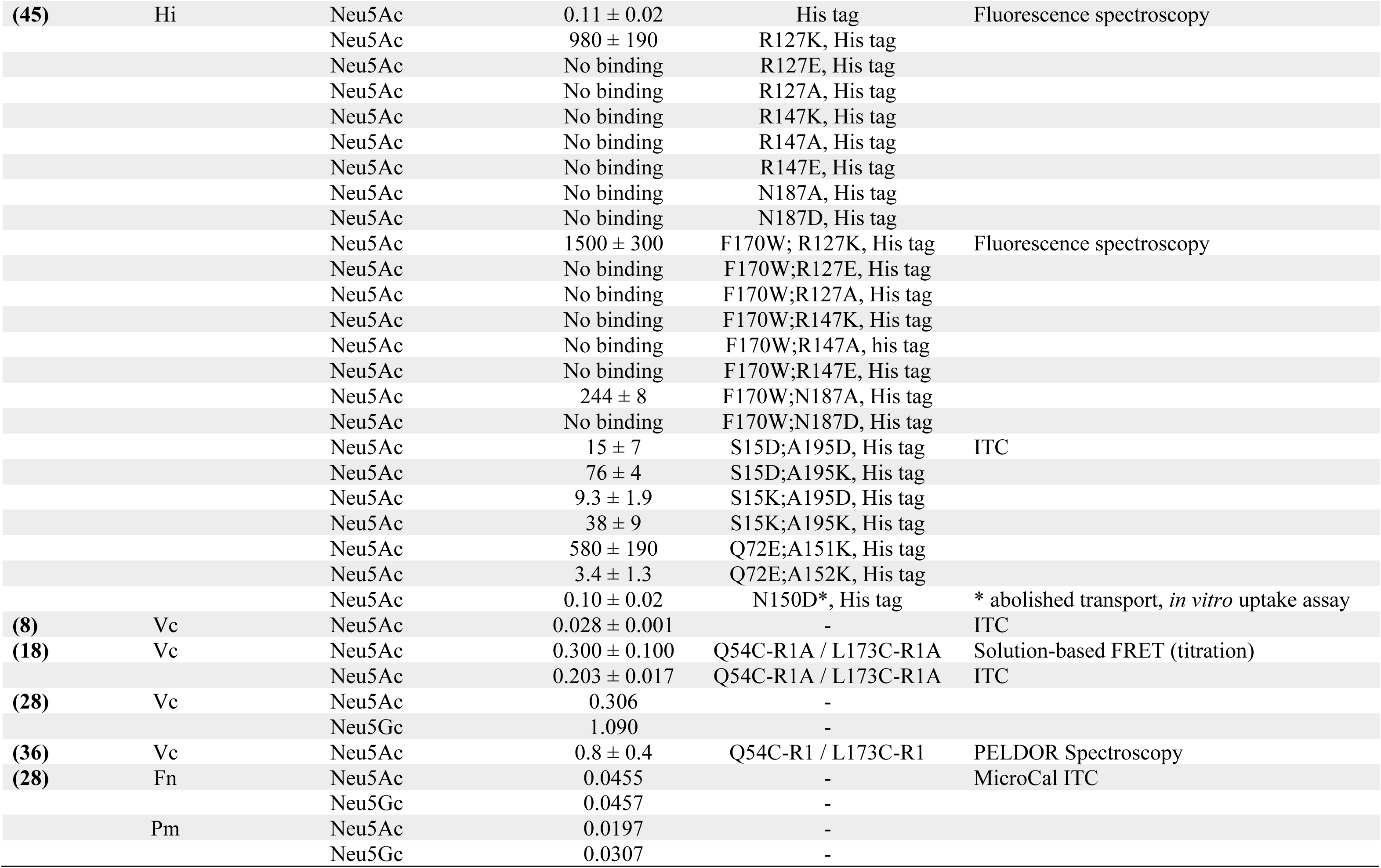
Summary of binding affinity data published for SiaP orthologues (as of April 24).

**Supplementary Table 2.**
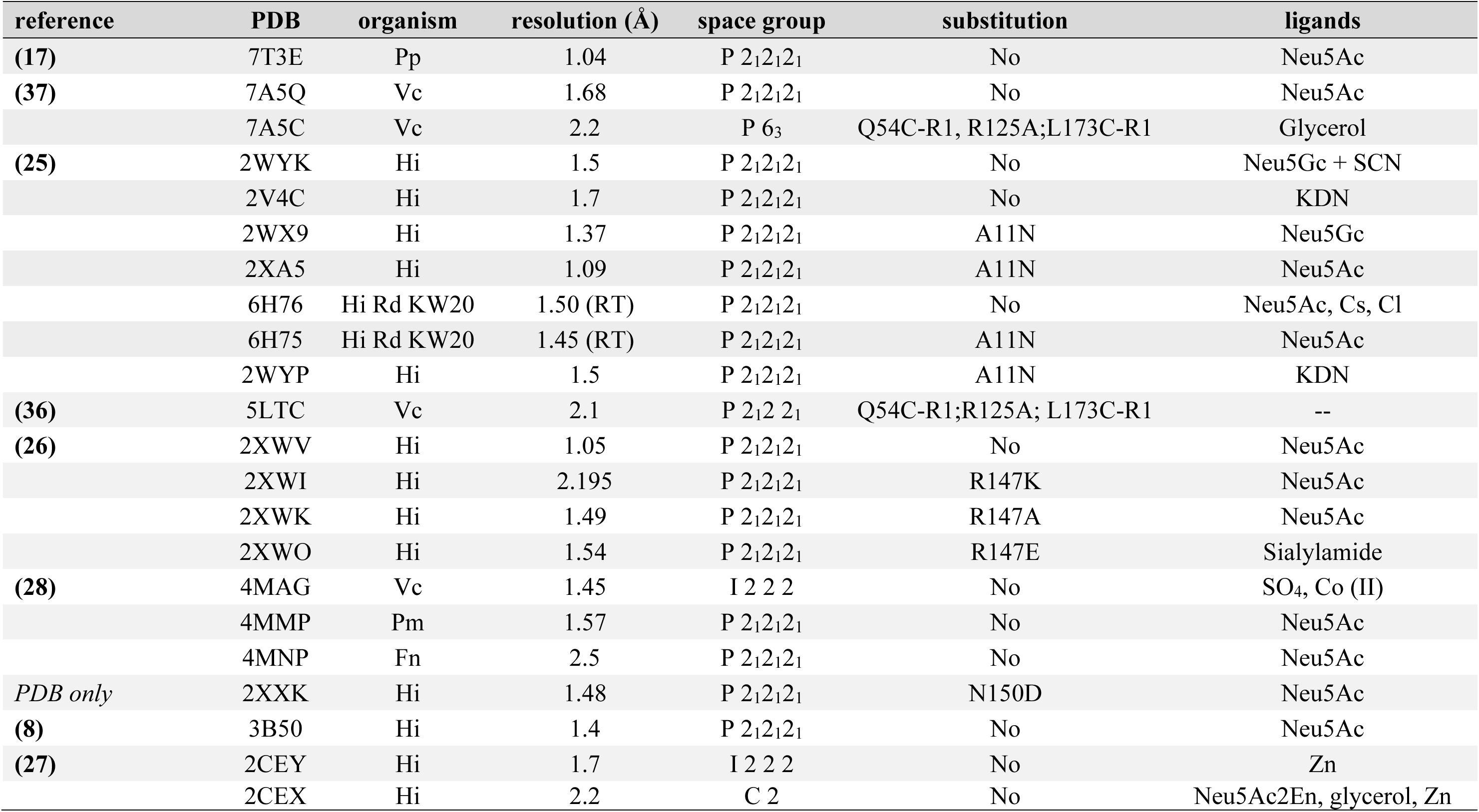
Summary of published SiaP structures on RCSB PDB (as of April 2024). Of the 22 structures deposited in the PDB, 15 are variation on *Hi*SiaP (mutants, different ligands) and four are *Vc*SiaP.

**Supplementary Table 3.** The root mean square difference (Å) between chains in the unbound *Aa*SiaP crystal asymmetric unit and the closed-bound structure, also compared with other SiaP homologues. Results show that the closed-unbound monomers of our structure (chains A and B) align best with the closed ligand-bound structure of *Aa*SiaP and homologous proteins *Pm*SiaP (75% sequence identity) and *Fn*SiaP (67% sequence identity).

|  |  | mostly closed-unbound |  | open-unbound |  | closed-bound |
| --- | --- | --- | --- | --- | --- | --- |
|  |  | chain A | chain B | chain C | chain D | + Neu5Ac |
| *Aa*SiaP | chain A mostly closed-unbound (9BH3) | - | 0.274 | 1.655 | 1.474 | 0.406 |
|  | chain B mostly closed-unbound (9BH3) | 0.274 | - | 1.620 | 1.541 | 0.285 |
|  | chain C open-unbound (9BH3) | 1.655 | 1.620 | - | 0.269 | 1.873 |
|  | chain D open-unbound (9BH3) | 1.474 | 1.541 | 0.269 | - | 1.722 |
|  | chain A + Neu5Ac (9BHF) | 0.406 | 0.285 | 1.873 | 1.722 | - |
| Other<br>SiaP's | *Hi*SiaP open-unbound (2CEY) | 1.759 | 1.683 | 0.471 | 0.469 | 1.876 |
|  | *Hi*SiaP closed-bound (6H76) | 0.515 | 0.415 | 2.097 | 1.914 | 0.440 |
|  | *Pm*SiaP closed-bound (4MMP) | 0.713 | 0.624 | 2.322 | 2.166 | 0.614 |
|  | *Fn*SiaP closed-bound (4MNP) | 0.765 | 0.630 | 2.004 | 1.843 | 0.640 |
|  | *Pp*SiaP closed-bound (7T3E) |  |  |  |  | 0.61 |
|  | *Vc*SiaP closed-bound (7A5Q) |  |  |  |  | 0.62 |
|  | *Vc*SiaP open-unbound (4MAG) | 2.530 | 2.521 | 1.221 | 1.308 | 2.599 |

**Supplementary Table 4.**
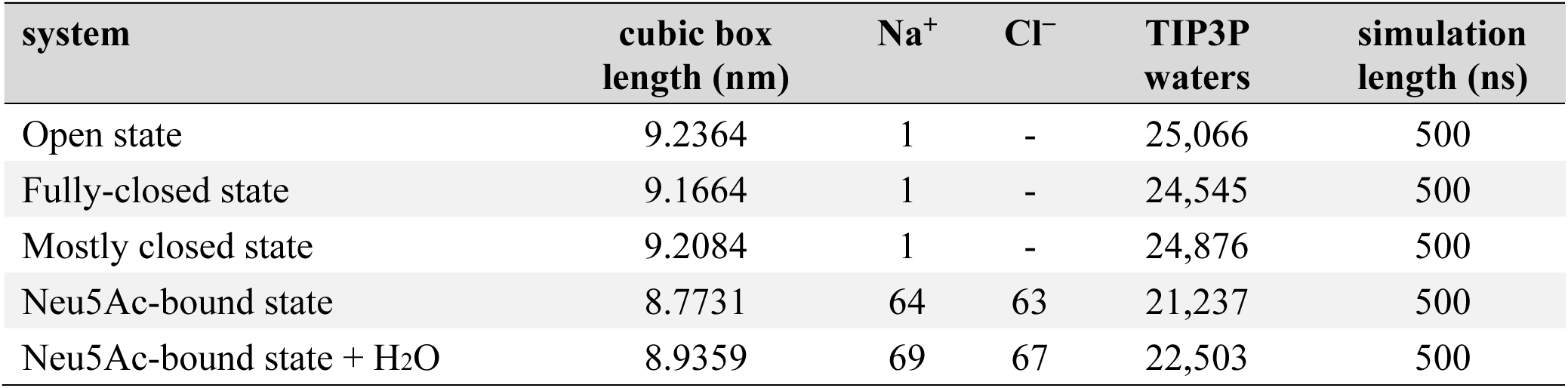
Cubic box length and total number of ions and water molecules in each system, and length of each simulation.

### Supplementary Figures

**Supplementary Figure 1.**
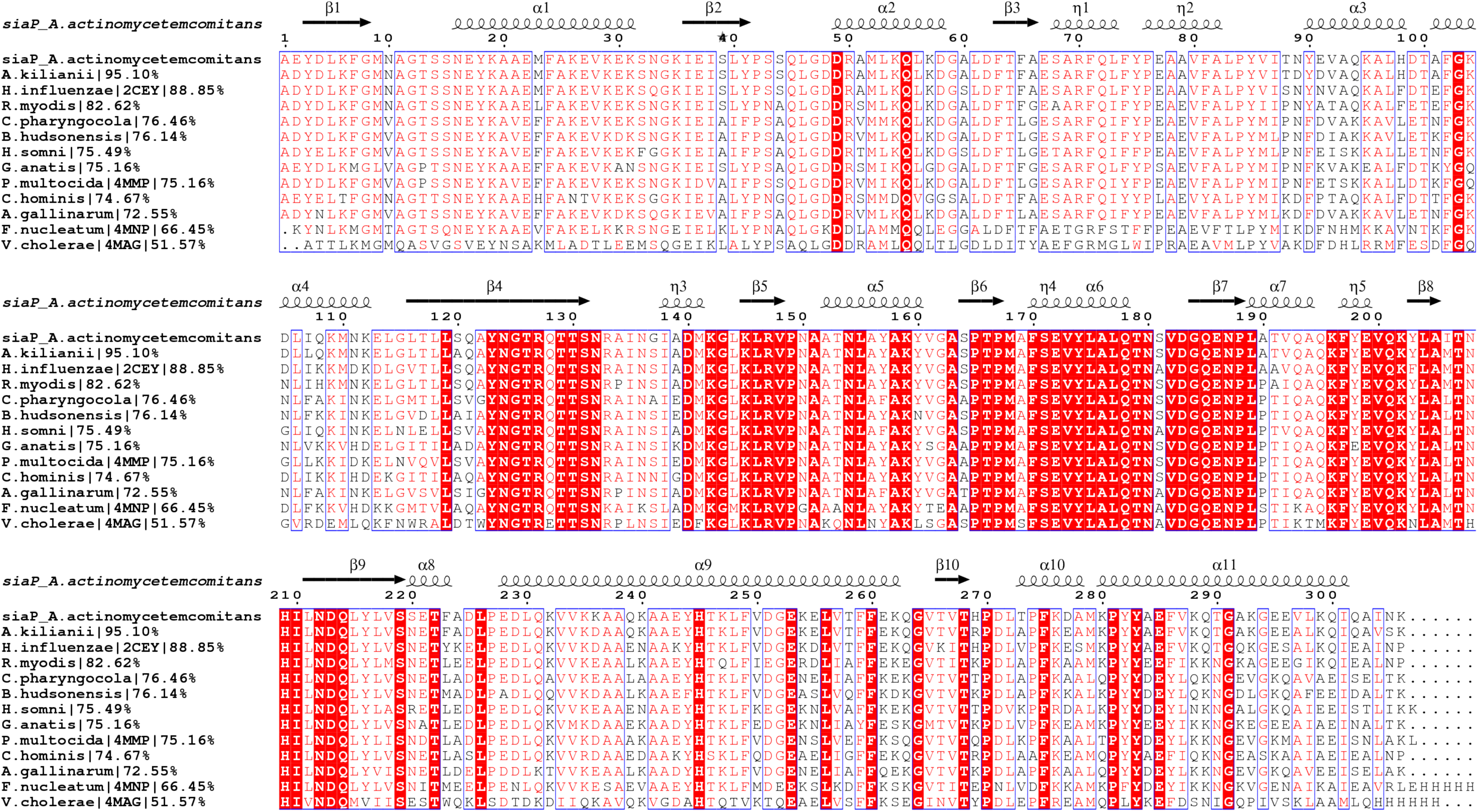
Multiple sequence alignment of non-redundant sequences of 12 *Aa*SiaP homologs (50–95% sequence identity), identified by PSI-BLAST using *Aa*SiaP gene as the query term.

**Supplementary Figure 2.**
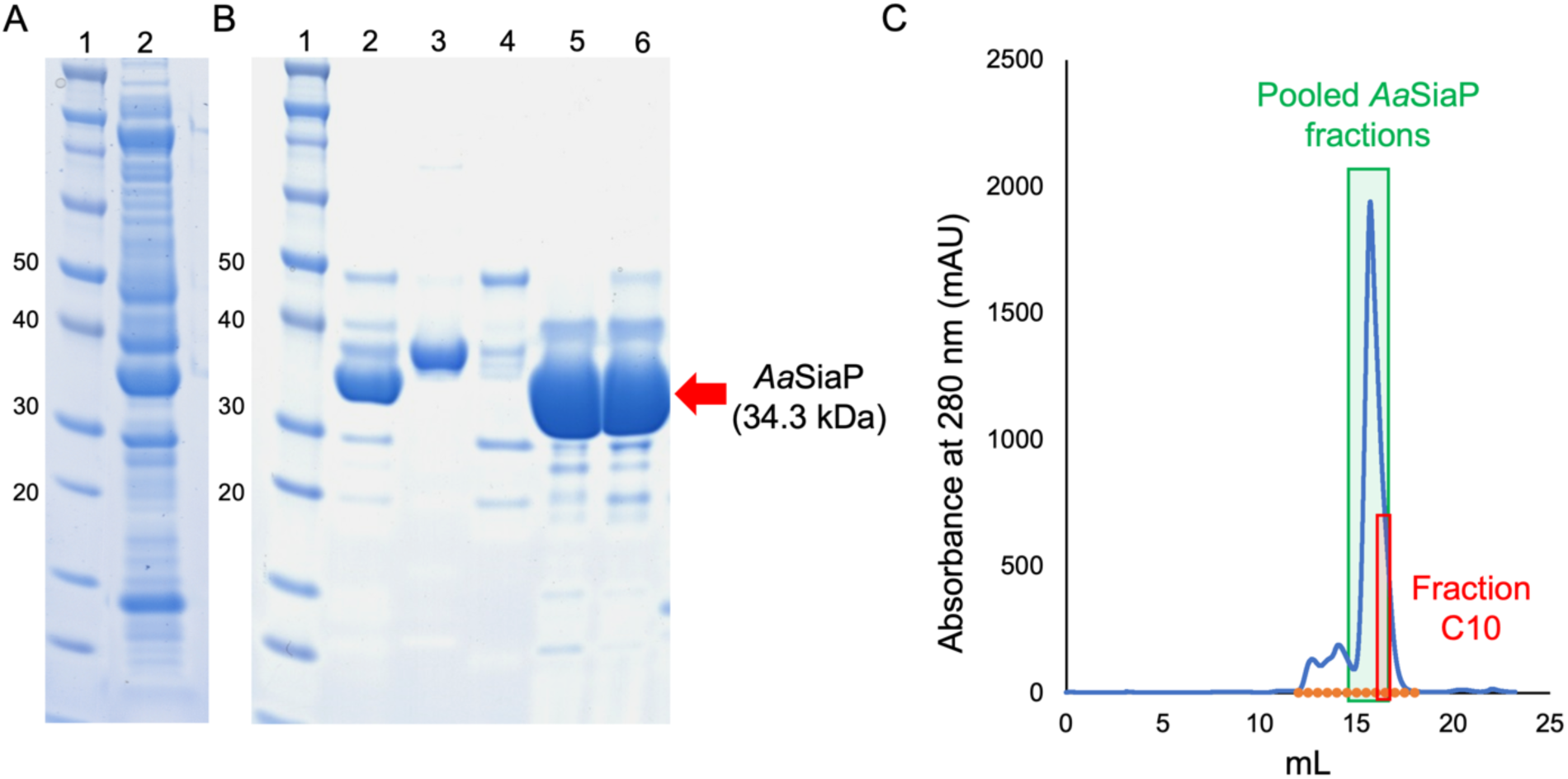
**A)** Recombinant *Aa*SiaP was highly expressed and could be isolated from the periplasmic fraction. Lane 1: molecular-weight ladder, lane 2: crude periplasmic lysate. **B)** Recombinant *Aa*SiaP was then purified by anion-exchange chromatography, hydrophobic-interaction chromatography, and size-exclusion chromatography. Lane 1: molecular-weight ladder, lane 2: before size-exclusion chromatography, lanes 3/4: flowthrough fraction from size-exclusion chromatography, lane 5: post size-exclusion chromatography *Aa*SiaP containing-fraction C10 (red box in C), lane 6: post size-exclusion chromatography *Aa*SiaP showing pooled fractions from 15– 16 mL (green box in C). **C)** *Aa*SiaP eluted from the SEC column as a single peak at approximately 15 mL, suggesting that it is a monodisperse species that remains stable throughout purification.

**Supplementary Figure 3.**
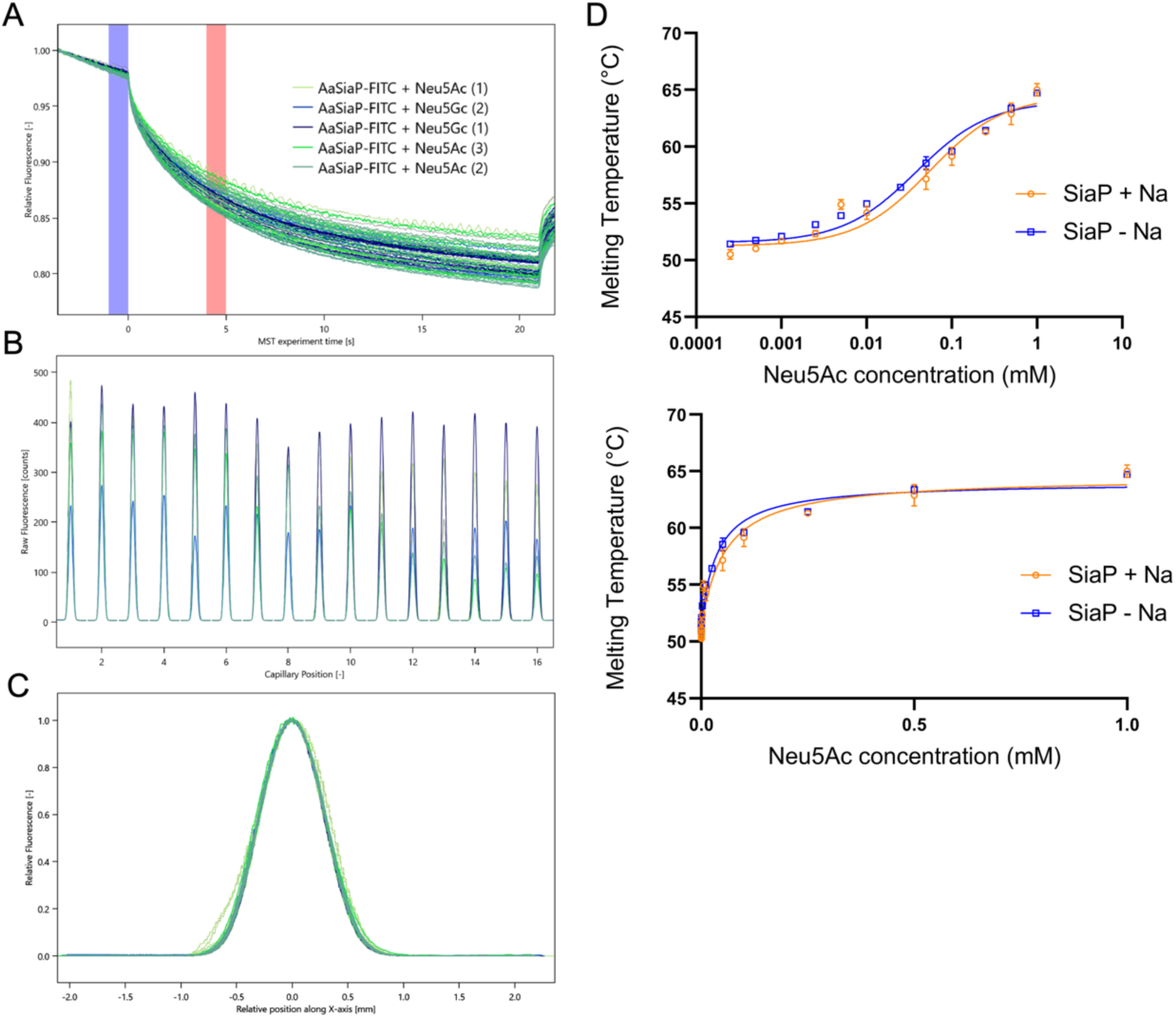
**A)** Raw MST traces for *Aa*SiaP-FITC with Neu5Ac (green curves) and Neu5Gc (blue curves), **B)** Capillary scans for MST, **C)** Capillary shape for MST data. **D)** Thermal shift assays with buffer containing-potassium salt instead of sodium had no effect on thermostability, suggesting that sodium is not required for binding. Titration data (as per Figure 2) plotted on a direct graph and fitted with a single-site ligand-binding model.

**Supplementary Figure 4.**
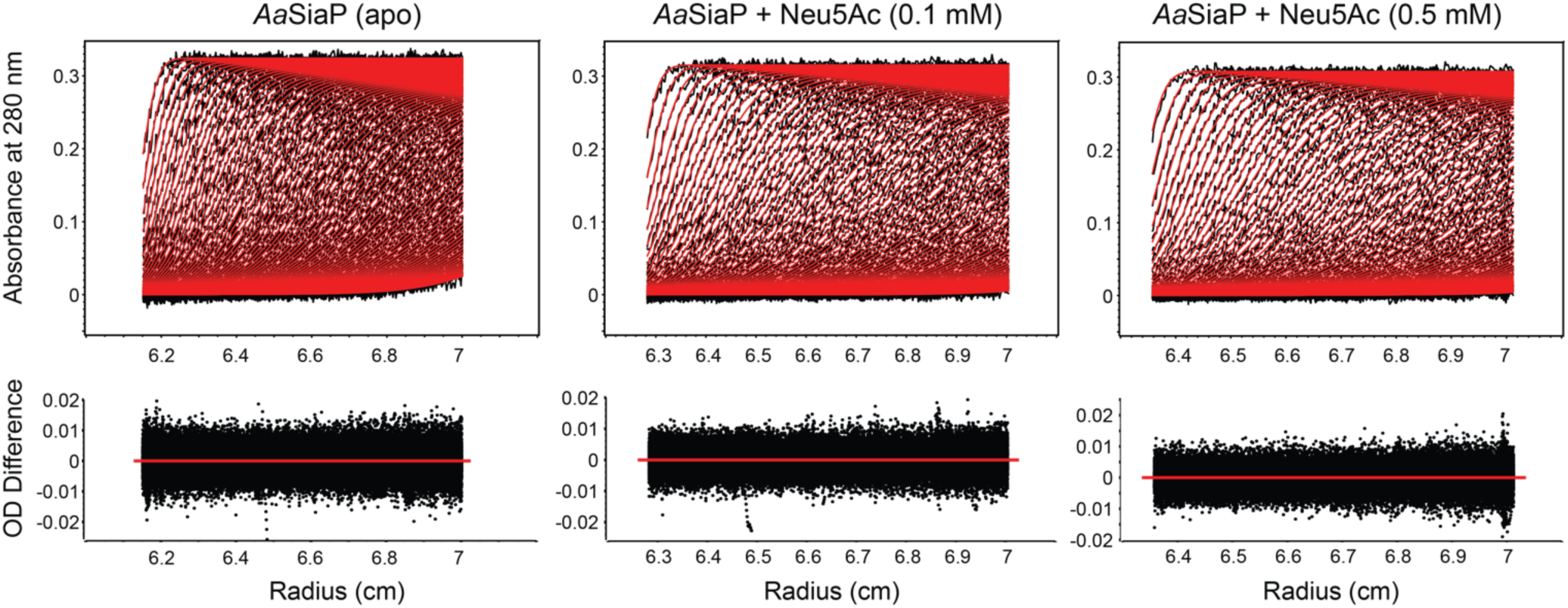
Sedimentation velocity data for *Aa*SiaP (black lines) and corresponding to the fit of the data to following 2DSA-Monte-Carlo solutions and parsimonious regularisation by genetic algorithm analysis (red lines) for the analysis shown in Figure 3B. The residuals for the fit are also displayed (bottom).

**Supplementary Figure 5.**
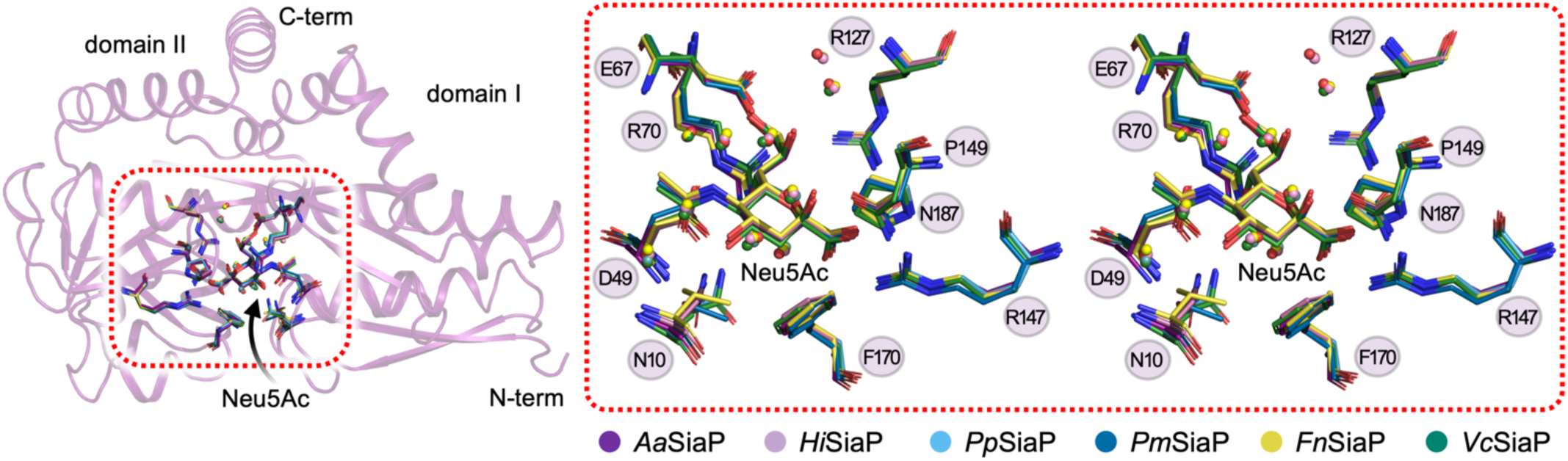
Neu5Ac bound to *Aa*SiaP and overlay with homologs. A focus on the Neu5Ac binding site demonstrates that both the contributing residues and the water shell around Neu5Ac is highly conserved. The inset is a cross-eyed stereo plot of the binding site.

**Supplementary Figure 6.**
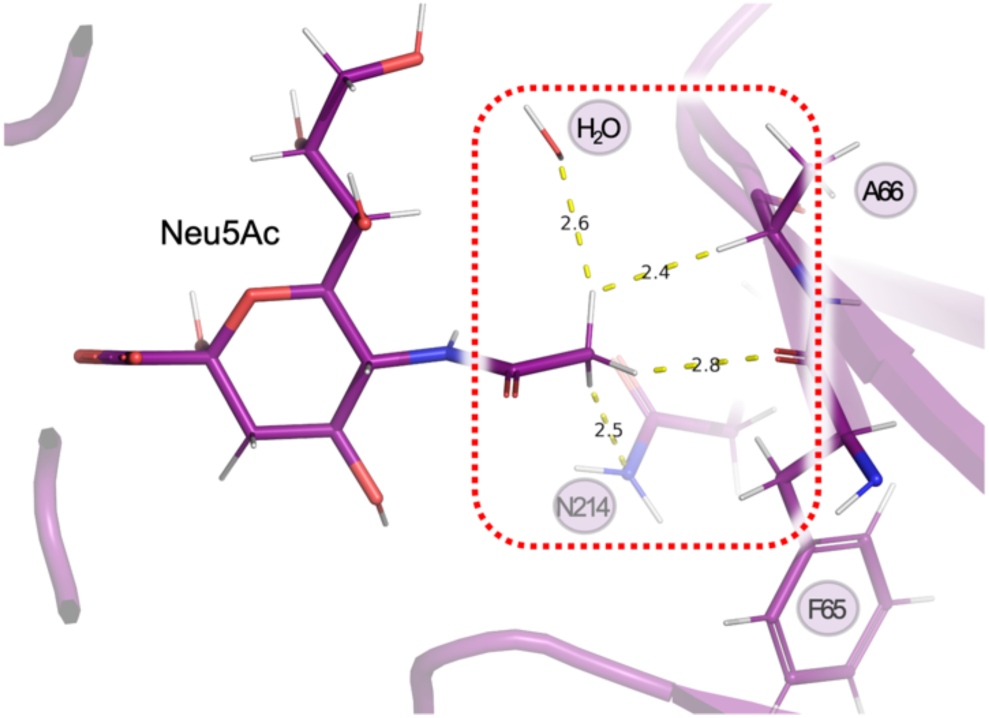
A focus on the binding interaction of the acetyl moiety of Neu5Ac.

## Notes

### Competing Interest Statement

The authors have declared no competing interest.

## References

1. Plumbridge, J., and Vimr, E. (1999) Convergent pathways for utilization of the amino sugars N-acetylglucosamine, N-acetylmannosamine, and N-acetylneuraminic acid by Escherichia coli. J. Bacteriol. 181, 47–54

2. Severi, E., Randle, G., Kivlin, P., Whitfield, K., Young, R., Moxon, R., Kelly, D., Hood, D., and Thomas, G. H. (2005) Sialic acid transport in Haemophilus influenzae is essential for lipopolysaccharide sialylation and serum resistance and is dependent on a novel tripartite ATP-independent periplasmic transporter. Mol Microbiol. 58, 1173–1185

3. Almagro-Moreno, S., and Boyd, E. F. (2009) Insights into the evolution of sialic acid catabolism among bacteria. BMC evolutionary biology. 9, 118

4. Thomas, G. H. (2016) Sialic acid acquisition in bacteria-one substrate, many transporters. Biochemical Society transactions. 44, 760–765

5. North, R. A., Horne, C. R., Davies, J. S., Remus, D. M., Muscroft-Taylor, A. C., Goyal, P., Wahlgren, W. Y., Ramaswamy, S., Friemann, R., and Dobson, R. C. J. (2018) “Just a spoonful of sugar…”: import of sialic acid across bacterial cell membranes. Biophysical Rev. 10, 219– 227

6. Chowdhury, N., Norris, J., McAlister, E., Lau, S. Y. K., Thomas, G. H., and Boyd, E. F. (2012) The VC1777–VC1779 proteins are members of a sialic acid-specific subfamily of TRAP transporters (SiaPQM) and constitute the sole route of sialic acid uptake in the human pathogen Vibrio cholerae. Microbiology+. 158, 2158–2167

7. Jenkins, G. A., Figueira, M., Kumar, G. A., Sweetman, W. A., Makepeace, K., Pelton, S. I., Moxon, R., and Hood, D. W. (2010) Sialic acid mediated transcriptional modulation of a highly conserved sialometabolism gene cluster in Haemophilus influenzae and its effect on virulence. BMC Microbiol. 10, 48

8. Johnston, J. W., Coussens, N. P., Allen, S., Houtman, J. C. D., Turner, K. H., Zaleski, A., Ramaswamy, S., Gibson, B. W., and Apicella, M. A. (2008) Characterization of the N-acetyl-5-neuraminic acid-binding site of the extracytoplasmic solute receptor (SiaP) of nontypeable Haemophilus influenzae strain 2019. J Biol Chem. 283, 855–865

9. Liang, Q., Ma, C., Crowley, S. M., Allaire, J. M., Han, X., Chong, R. W. W., Packer, N. H., Yu, H. B., and Vallance, B. A. (2023) Sialic acid plays a pivotal role in licensing Citrobacter rodentium’s transition from the intestinal lumen to a mucosal adherent niche. Proc. Natl. Acad. Sci. United States Am. 120, e2301115120

10. Coombes, D., Davies, J. S., Newton-Vesty, M. C., Horne, C. R., Setty, T. G., Subramanian, R., Moir, J. W. B., Friemann, R., Panjikar, S., Griffin, M. D. W., North, R. A., and Dobson, R. C. J. (2020) The basis for non-canonical ROK family function in the N-acetylmannosamine kinase from the pathogen Staphylococcus aureus. J Biol Chem. 295, 3301–3315

11. Currie, M. J., Manjunath, L., Horne, C. R., Rendle, P. M., Subramanian, R., Friemann, R., Fairbanks, A. J., Muscroft-Taylor, A. C., North, R. A., and Dobson, R. C. J. (2021) N-Acetylmannosamine-6-phosphate 2-epimerase uses a novel substrate-assisted mechanism to catalyze amino sugar epimerization. J Biol Chem. 297, 101113

12. Setty, T. G., Sarkar, A., Coombes, D., Dobson, R. C. J., and Subramanian, R. (2020) Structure and function of N-acetylmannosamine kinases from pathogenic bacteria. Acs Omega. 5, 30923–30936

13. Horne, C. R., Kind, L., Davies, J. S., and Dobson, R. C. J. (2020) On the structure and function of Escherichia coli YjhC: An oxidoreductase involved in bacterial sialic acid metabolism. Proteins Struct Funct Bioinform. 88, 654–668

14. North, R. A., Watson, A. J. A., Pearce, F. G., Muscroft-Taylor, A. C., Friemann, R., Fairbanks, A. J., and Dobson, R. C. J. (2016) Structure and inhibition of N-acetylneuraminate lyase from methicillin-resistant Staphylococcus aureus. FEBS Letters. 590, 4414–4428

15. Kelly, D. J., and Thomas, G. H. (2001) The tripartite ATP-independent periplasmic (TRAP) transporters of bacteria and archaea. Fems Microbiol Rev. 25, 405–424

16. Mulligan, C., Fischer, M., and Thomas, G. H. (2011) Tripartite ATP-independent periplasmic (TRAP) transporters in bacteria and archaea. Fems Microbiol Rev. 35, 68–86

17. Davies, J. S., Currie, M. J., North, R. A., Scalise, M., Wright, J. D., Copping, J. M., Remus, D. M., Gulati, A., Morado, D. R., Jamieson, S. A., Newton-Vesty, M. C., Abeysekera, G. S., Ramaswamy, S., Friemann, R., Wakatsuki, S., Allison, J. R., Indiveri, C., Drew, D., Mace, P. D., and Dobson, R. C. J. (2023) Structure and mechanism of a tripartite ATP-independent periplasmic TRAP transporter. Nat Commun. 14, 1120

18. Peter, M. F., Ruland, J. A., Depping, P., Schneberger, N., Severi, E., Moecking, J., Gatterdam, K., Tindall, S., Durand, A., Heinz, V., Siebrasse, J. P., Koenig, P.-A., Geyer, M., Ziegler, C., Kubitscheck, U., Thomas, G. H., and Hagelueken, G. (2022) Structural and mechanistic analysis of a tripartite ATP-independent periplasmic TRAP transporter. Nat Commun. 13, 4471

19. Currie, M. J., Davies, J. S., Scalise, M., Gulati, A., Wright, J. D., Newton-Vesty, M. C., Abeysekera, G. S., Subramanian, R., Wahlgren, W. Y., Friemann, R., Allison, J. R., Mace, P. D., Griffin, M. D. W., Demeler, B., Wakatsuki, S., Drew, D., Indiveri, C., Dobson, R. C. J., and North, R. A. (2023) Structural and biophysical analysis of a Haemophilus influenzae tripartite ATP-independent periplasmic (TRAP) transporter. eLife

20. Goyal, P., Dhanabalan, K. V., Scalise, M., Friemann, R., Indiveri, C., Dobson, R. C. J., Vinothkumar, K. R., and Ramaswamy, S. (2024) Molecular determinants of Neu5Ac binding to a tripartite ATP independent periplasmic (TRAP) transporter. bioRxiv. 10.1101/2024.03.29.587382

21. Davies, J. S., Currie, M. J., Dobson, R. C. J., Horne, C. R., and North, R. A. (2023) TRAPs: the ‘elevator-with-an-operator’ mechanism. Trends in Biochemical Sciences

22. Maqbool, A., Horler, R. S. P., Muller, A., Wilkinson, A. J., Wilson, K. S., and Thomas, G. H. (2015) The substrate-binding protein in bacterial ABC transporters: dissecting roles in the evolution of substrate specificity. Biochem. Soc. Trans. 43, 1011–1017

23. Fischer, M., Zhang, Q. Y., Hubbard, R. E., and Thomas, G. H. (2010) Caught in a TRAP: substrate-binding proteins in secondary transport. Trends Microbiol. 18, 471–8

24. Mulligan, C., Geertsma, E. R., Severi, E., Kelly, D. J., Poolman, B., and Thomas, G. H. (2009) The substrate-binding protein imposes directionality on an electrochemical sodium gradient-driven TRAP transporter. Proc National Acad Sci USA. 106, 1778–1783

25. Darby, J. F., Hopkins, A. P., Shimizu, S., Roberts, S. M., Brannigan, J. A., Turkenburg, J. P., Thomas, G. H., Hubbard, R. E., and Fischer, M. (2019) Water Networks Can Determine the Affinity of Ligand Binding to Proteins. J Am Chem Soc. 141, 15818–15826

26. Fischer, M., Hopkins, A. P., Severi, E., Hawkhead, J., Bawdon, D., Watts, A. G., Hubbard, R. E., and Thomas, G. H. (2015) Tripartite ATP-independent Periplasmic (TRAP) Transporters Use an Arginine-mediated Selectivity Filter for High Affinity Substrate Binding*. J Biol Chem. 290, 27113–27123

27. Müller, A., Severi, E., Mulligan, C., Watts, A. G., Kelly, D. J., Wilson, K. S., Wilkinson, A. J., and Thomas, G. H. (2006) Conservation of structure and mechanism in primary and secondary transporters exemplified by SiaP, a sialic acid binding virulence factor from Haemophilus influenzae. J Biol Chem. 281, 22212–22222

28. Setty, T. G., Cho, C., Govindappa, S., Apicella, M. A., and Ramaswamy, S. (2014) Bacterial periplasmic sialic acid-binding proteins exhibit a conserved binding site. Acta Crystallogr D. 70, 1801–1811

29. Berntsson, R. P. A., Smits, S. H. J., Schmitt, L., Slotboom, D.-J., and Poolman, B. (2010) A structural classification of substrate-binding proteins. FEBS Lett. 584, 2606–2617

30. Mao, B., Pear, M. R., McCammon, J. A., and Quiocho, F. A. (1982) Hinge-bending in L-arabinose-binding protein. The “Venus’s-flytrap” model. J. Biol. Chem. 257, 1131–1133

31. Felder, C. B., Graul, R. C., Lee, A. Y., Merkle, H. P., and Sadee, W. (1999) The Venus flytrap of periplasmic binding proteins: an ancient protein module present in multiple drug receptors. AAPS pharmSci. 1, E2

32. Flocco, M. M., and Mowbray, S. L. (1994) The 1.9 A x-ray structure of a closed unliganded form of the periplasmic glucose/galactose receptor from Salmonella typhimurium. J. Biol. Chem. 269, 8931–6

33. Zhang, Y., Mannering, D. E., Davidson, A. L., Yao, N., and Manson, M. D. (1996) Maltose-binding Protein Containing an Interdomain Disulfide Bridge Confers a Dominant-negative Phenotype for Transport and Chemotaxis. J. Biol. Chem. 271, 17881–17889

34. Tang, C., Schwieters, C. D., and Clore, G. M. (2007) Open-to-closed transition in apo maltose-binding protein observed by paramagnetic NMR. Nature. 449, 1078–1082

35. Oswald, C., Smits, S. H. J., Höing, M., Sohn-Bösser, L., Dupont, L., Rudulier, D. L., Schmitt, L., and Bremer, E. (2008) Crystal Structures of the Choline/Acetylcholine Substrate-binding Protein ChoX from Sinorhizobium meliloti in the Liganded and Unliganded-Closed States*. J. Biol. Chem. 283, 32848–32859

36. Glaenzer, J., Peter, M. F., Thomas, G. H., and Hagelueken, G. (2017) PELDOR spectroscopy reveals two defined states of a sialic acid TRAP transporter SBP in solution. Biophysical journal. 112, 109–120

37. Peter, M. F., Gebhardt, C., Glaenzer, J., Schneberger, N., Boer, M. de, Thomas, G. H., Cordes, T., and Hagelueken, G. (2020) Triggering closure of a sialic acid TRAP transporter substrate binding protein through binding of natural or artificial substrates. J Mol Biol. 433, 166756

38. Marinelli, F., and Fiorin, G. (2019) Structural Characterization of Biomolecules through Atomistic Simulations Guided by DEER Measurements. Structure. 27, 359–370.e12

39. Winkelhoff, A. J., and Slots, J. (1999) Actinobacillus actinomycetemcomitans and Porphyromonas gingivalis in nonoral infections. Periodontol. 2000. 20, 122–135

40. Gholizadeh, P., Pormohammad, A., Eslami, H., Shokouhi, B., Fakhrzadeh, V., and Kafil, H. S. (2017) Oral pathogenesis of Aggregatibacter actinomycetemcomitans. Microb. Pathog. 113, 303–311

41. Maeda, H., and Kawauchi, H. (1968) A new method for the determination of N-terminus of peptides chain with fluorescein-isothiocyanate. Biochem. Biophys. Res. Commun. 31, 188–192

42. Williams, C. J., Headd, J. J., Moriarty, N. W., Prisant, M. G., Videau, L. L., Deis, L. N., Verma, V., Keedy, D. A., Hintze, B. J., Chen, V. B., Jain, S., Lewis, S. M., Arendall, W. B., Snoeyink, J., Adams, P. D., Lovell, S. C., Richardson, J. S., and Richardson, D. C. (2018) MolProbity: More and better reference data for improved all-atom structure validation. Protein Sci. 27, 293– 315

43. Chen, V. B., Arendall, W. B., Headd, J. J., Keedy, D. A., Immormino, R. M., Kapral, G. J., Murray, L. W., Richardson, J. S., and Richardson, D. C. (2010) MolProbity: all-atom structure validation for macromolecular crystallography. Acta Crystallogr Sect D Biological Crystallogr. 66, 12–21

44. Krissinel, E., and Henrick, K. (2007) Inference of macromolecular assemblies from crystalline state. J Mol Bio. 372, 774–797

45. Hopkins, A. P. (2010) Molecular and Biochemical Characterisation of SiaP as a Sialic Acid Binding Protein Component of a TRAP Transporter of Sialic Acid. Ph.D. thesis, The University of York

46. Skou, S., Gillilan, R. E., and Ando, N. (2014) Synchrotron-based small-angle X-ray scattering of proteins in solution. Nat. Protoc. 9, 1727–1739

47. Boer, M. de, Gouridis, G., Vietrov, R., Begg, S. L., Schuurman-Wolters, G. K., Husada, F., Eleftheriadis, N., Poolman, B., McDevitt, C. A., and Cordes, T. (2019) Conformational and dynamic plasticity in substrate-binding proteins underlies selective transport in ABC importers. eLife. 8, e44652

48. Fiorin, G., Marinelli, F., and Faraldo-Gómez, J. D. (2019) Direct Derivation of Free Energies of Membrane Deformation and Other Solvent Density Variations From Enhanced Sampling Molecular Dynamics. J. Comput. Chem. 41, 449–459

49. Peter, M. F., Ruland, J. A., Kim, Y., Hendricks, P., Schneberger, N., Siebrasse, J. P., Thomas, G. H., Kubitscheck, U., and Hagelueken, G. (2024) Conformational coupling of the sialic acid TRAP transporter HiSiaQM with its substrate binding protein HiSiaP. Nat. Commun. 15, 217

50. Oldham, M. L., Khare, D., Quiocho, F. A., Davidson, A. L., and Chen, J. (2007) Crystal structure of a catalytic intermediate of the maltose transporter. Nature. 450, 515–521

51. Gouridis, G., Schuurman-Wolters, G. K., Ploetz, E., Husada, F., Vietrov, R., Boer, M. de, Cordes, T., and Poolman, B. (2014) Conformational dynamics in substrate-binding domains influences transport in the ABC importer GlnPQ. Nat. Struct. Mol. Biol. 22, 57–64

52. Madeira, F., Park, Y. mi, Lee, J., Buso, N., Gur, T., Madhusoodanan, N., Basutkar, P., Tivey, A. R. N., Potter, S. C., Finn, R. D., and Lopez, R. (2019) The EMBL-EBI search and sequence analysis tools APIs in 2019. Nucleic Acids Res. 47, W636–W641

53. Robert, X., and Gouet, P. (2014) Deciphering key features in protein structures with the new ENDscript server. Nucleic Acids Research. 42, W320–4

54. Armenteros, J. J. A., Tsirigos, K. D., Sønderby, C. K., Petersen, T. N., Winther, O., Brunak, S., Heijne, G. von, and Nielsen, H. (2019) SignalP 5.0 improves signal peptide predictions using deep neural networks. Nat. Biotechnol. 37, 420–423

55. Vivoli, M., Novak, H. R., Littlechild, J. A., and Harmer, N. J. (2014) Determination of Protein-ligand Interactions Using Differential Scanning Fluorimetry. J. Vis. Exp. 10.3791/51809-v

56. Demeler, B. (2010) Methods for the design and analysis of sedimentation velocity and sedimentation equilibrium experiments with proteins. *Current protocols in protein science / editorial board, John E. Coligan … [et al.]*. **Chapter 7**, Unit 7.13

57. Demeler, B. (2005) UltraScan - A comprehensive data analysis software package for analytical ultracentrifugation experiments. in *Modern Analytical Ultracentrifugation: Techniques and Methods*. (Scott, D. J., Rowe., S. E. H. and A. J., and Rowe., A. J. eds)

58. Brookes, E., Cao, W., and Demeler, B. (2010) A two-dimensional spectrum analysis for sedimentation velocity experiments of mixtures with heterogeneity in molecular weight and shape. European biophysics journal: EBJ. 39, 405–414

59. Brookes, E. H., and Demeler, B. (2007) Parsimonious regularization using genetic algorithms applied to the analysis of analytical ultracentrifugation experiments, pp. 361–368, the 9th annual conference, 10.1145/1276958.1277035

60. Fleming, P. J., and Fleming, K. G. (2018) HullRad: Fast Calculations of Folded and Disordered Protein and Nucleic Acid Hydrodynamic Properties. Biophys. J. 114, 856–869

61. Kabsch, W. (2010) XDS. Acta Crystallogr. D. 66, 125–132

62. Potterton, L., Agirre, J., Ballard, C., Cowtan, K., Dodson, E., Evans, P. R., Jenkins, H. T., Keegan, R., Krissinel, E., Stevenson, K., Lebedev, A., McNicholas, S. J., Nicholls, R. A., Noble, M., Pannu, N. S., Roth, C., Sheldrick, G., Skubak, P., Turkenburg, J., Uski, V., Delft, F. von, Waterman, D., Wilson, K., Winn, M., and Wojdyr, M. (2018) CCP4i2: the new graphical user interface to the CCP4 program suite. Acta Crystallogr Sect D. 74, 68–84

63. McCoy, A. J., Grosse-Kunstleve, R. W., Adams, P. D., Winn, M. D., Storoni, L. C., and Read, R. J. (2007) Phaser crystallographic software. J Appl Crystallogr. 40, 658–674

64. Emsley, P., and Cowtan, K. (2004) Coot: model-building tools for molecular graphics. Acta Crystallogr Sect D Biological Crystallogr. 60, 2126–2132

65. Murshudov, G. N., Vagin, A. A., and Dodson, E. J. (1997) Refinement of macromolecular structures by the maximum-likelihood method. Acta Crystallogr Sect D Biological Crystallogr. 53, 240–255

66. Murshudov, G. N., Skubák, P., Lebedev, A. A., Pannu, N. S., Steiner, R. A., Nicholls, R. A., Winn, M. D., Long, F., and Vagin, A. A. (2011) REFMAC5 for the refinement of macromolecular crystal structures. Acta Crystallogr. Sect. D: Biol. Crystallogr. 67, 355–367

67. Liebschner, D., Afonine, P. V., Baker, M. L., Bunkóczi, G., Chen, V. B., Croll, T. I., Hintze, B., Hung, L.-W., Jain, S., McCoy, A. J., Moriarty, N. W., Oeffner, R. D., Poon, B. K., Prisant, M. G., Read, R. J., Richardson, J. S., Richardson, D. C., Sammito, M. D., Sobolev, O. V., Stockwell, D. H., Terwilliger, T. C., Urzhumtsev, A. G., Videau, L. L., Williams, C. J., and Adams, P. D. (2019) Macromolecular structure determination using X-rays, neutrons and electrons: recent developments in Phenix. Acta Crystallogr D. 75, 861–877

68. Emsley, P., Lohkamp, B., Scott, W. G., and Cowtan, K. (2010) Features and development of Coot. Acta Crystallographica Section D Biological Crystallography. 66, 486–501

69. Ryan, T. M., Trewhella, J., Murphy, J. M., Keown, J. R., Casey, L., Pearce, F. G., Goldstone, D. C., Chen, K., Luo, Z., Kobe, B., McDevitt, C. A., Watkin, S. A., Hawley, A. M., Mudie, S. T., Boban, V. S., and Kirby, N. (2018) An optimized SEC-SAXS system enabling high X-ray dose for rapid SAXS assessment with correlated UV measurements for biomolecular structure analysis. Journal of applied crystallography. 51, 97–111

70. Franke, D., Petoukhov, M. V., Konarev, P. V., Panjkovich, A., Tuukkanen, A., Mertens, H. D. T., Kikhney, A. G., Hajizadeh, N. R., Franklin, J. M., Jeffries, C. M., and Svergun, D. I. (2017) ATSAS 2.8: a comprehensive data analysis suite for small-angle scattering from macromolecular solutions. J Appl Crystallogr. 50, 1212–1225

71. Svergun, D., Barberato, C., and Koch, M. (1995) CRYSOL – a program to evaluate X-ray solution scattering of biological macromolecules from atomic coordinates. J Appl Crystallogr. 28, 768–773

72. Petoukhov, M. V., and Svergun, D. I. (2015) Ambiguity assessment of small-angle scattering curves from monodisperse systems. Acta Crystallogr. Sect. D: Biol. Crystallogr. 71, 1051– 1058

73. Svergun, D. I., Petoukhov, M. V., and Koch, M. H. J. (2001) Determination of Domain Structure of Proteins from X-Ray Solution Scattering. Biophys. J. 80, 2946–2953

74. Volkov, V. V., and Svergun, D. I. (2003) Uniqueness of ab initio shape determination in small-angle scattering. J. Appl. Crystallogr. 36, 860–864

75. Panjkovich, A., and Svergun, D. I. (2016) Deciphering conformational transitions of proteins by small angle X-ray scattering and normal mode analysis. Physical Chemistry Chemical Physics. 18, 5707–5719

76. Abraham, M. J., Murtola, T., Schulz, R., Páll, S., Smith, J. C., Hess, B., and Lindahl, E. (2015) GROMACS: High performance molecular simulations through multi-level parallelism from laptops to supercomputers. SoftwareX. 1–2, 19–25

77. Huang, J., and MacKerell, A. D. (2013) CHARMM36 all-atom additive protein force field: validation based on comparison to NMR data. Journal of computational chemistry. 34, 2135– 2145

78. Darden, T., York, D., and Pedersen, L. (1993) Particle mesh Ewald: An N ⋅log(N) method for Ewald sums in large systems. J. Chem. Phys. 98, 10089–10092

79. Verlet, L. (1967) Computer “Experiments” on Classical Fluids. I. Thermodynamical Properties of Lennard-Jones Molecules. Phys. Rev. 159, 98–103

80. Hess, B., Bekker, H., Berendsen, H. J. C., and Fraaije, J. G. E. M. (1997) LINCS: A linear constraint solver for molecular simulations. J. Comput. Chem. 18, 1463–1472

81. Bussi, G., Donadio, D., and Parrinello, M. (2007) Canonical sampling through velocity rescaling. J. Chem. Phys. 10.1063/1.2408420

82. Berendsen, H. J. C., Postma, J. P. M., Gunsteren, W. F. van, DiNola, A., and Haak, J. R. (1984) Molecular dynamics with coupling to an external bath. J. Chem. Phys. 81, 3684–3690

83. Parrinello, M., and Rahman, A. (1980) Crystal Structure and Pair Potentials: A Molecular-Dynamics Study. Phys. Rev. Lett. 45, 1196–1199

84. Jorgensen, W. L., Chandrasekhar, J., Madura, J. D., Impey, R. W., and Klein, M. L. (1983) Comparison of simple potential functions for simulating liquid water. J. Chem. Phys. 79, 926– 935

85. Humphrey, W., Dalke, A., and Schulten, K. (1996) VMD: Visual molecular dynamics. J. Mol. Graph. 14, 33–38

86. Team, R. C. (2021) R: A language and environment for statistical computing, Vienna, Austria.

